# Impaired astrocytic Cx43 syncytial network in lateral habenula affects synaptic connectivity and drives adolescent stress-induced depression

**DOI:** 10.64898/2026.10.02.756205

**Authors:** Hui Li, Qi Wang, Yixun Su, Hui Chen, Zhonghao Wu, Xiaoying Chen, Chenmeng Wang, Yafei Zhao, Dan Li, Xiaomin Huang, Rao Fu, Wenhui Huang, Yong Tang, Jianqin Niu, Alexei Verkhratsky, Chenju Yi

## Abstract

Adolescent depression is a major neuropsychiatric disorder induced by early life stress; the underlying cellular mechanisms instigating pathological circuit malformation remain unclear, and, consequently, effective therapeutic strategy is limited. Here, we analysed the contribution of astrocytes to early life stress-induced adolescent depression using a mouse model of parental separation. We identified a previously unrecognised regional specialisation of lateral habenula (LHb) astrocytes, characterised by compact cellular domains, abundant leaflets, and enhanced Cx43 gap junctional coupling. Parental separation selectively reduced astrocytic Cx43 expression and gap junctional communication in the LHb, which was accompanied by profound astrocytic atrophy and loss of leaflets. Single-cell sequencing further revealed that parental separation induces a broad reorganisation of LHb astrocyte transcriptional states, including metabolic, stress-response, and inflammatory programs; moreover, it specifically reduces astrocyte-neuron communication. Consistent with this, we found that structural remodelling of LHb astrocytes affected tripartite synapse geometry, resulting in impaired excitatory and inhibitory synaptic transmission, and neuronal Ca^2+^ hyperexcitability, all ultimately associated with depressive-like behaviours. Selective knockdown of astrocytic Cx43 in the LHb recapitulated these pathological changes and translated into depressive-like behaviours. Mechanistically, we found that impaired Cx43 gap junctional communication is associated with excessive astrocytic Ca^2+^ dynamics, resulting in Rac1-GTPase activation and decreased p-Ezrin levels. *In vivo* inhibition of LHb astrocytic Ca^2+^ signalling rescued astrocytic atrophy and depressive-like behaviours following parental separation. Together, our findings establish a Cx43-Ca^2+^-Rac1-p-Ezrin pathway that links gap junction malfunction to structural remodelling of astrocytes and disruption of neuron–astrocyte interactions in the developing LHb. Our study reveals astrocytic Ca^2+^ dysregulation as an active downstream effector of Cx43 dysfunction and identifies a regionally specialised astrocytic mechanism through which early-life stress disrupts synaptic transmission and neuronal function during adolescence, providing a potential cellular target for intervention in adolescent depression.

## Introduction

Childhood and adolescent depression have emerged as a global challenge to public health. Early-life stress, such as separation from parents, is known to disrupt the function of multiple brain regions^1–3^, thereby increasing the susceptibility to depression in childhood and adolescence^4, 5^. However, contemporary antidepressants, exclusively developed under a neuron□centric paradigm and primarily targeting monoaminergic synapses, show limited efficacy in youth ^4,5^. The limited efficacy of antidepressants exposes a major gap in the field: neuronal mechanisms alone are unlikely to fully explain pathophysiology of depression. This is particularly important for adolescent depression, where ongoing neurodevelopment may shape disease processes that differ fundamentally from those in adults.

Astrocytes are widely recognised as active regulator of synaptic transmission and plasticity through their close association with neuronal synapses, i.e., tripartite synapses ^6–8, 9^, whereas astrocyte malfunction has been increasingly implicated in psychiatric disorders ^10,11^. In particular, depression in humans and depressive-like behaviours in animal models are associated with marked atrophy and aberrant function of astrocytes in the medial prefrontal cortex (mPFC) and amygdala, supporting a pathogenetic role for astrocytes beyond neuronal dysfunction ^12–15^. However, these findings are largely derived from the adult animals. Childhood and adolescence are critical periods of synaptic pruning and circuit refinement ^16^, yet whether astrocytes shape the vulnerability to depression during this developmental window remains poorly understood. In addition, astrocytes show marked regional heterogeneity in morphology, gene expression and function^17,18^, suggesting that their contribution to diseases must be resolved in a region-specific manner. The lateral habenula (LHb), a core anti-reward hub implicated in depression, provides a key site to this question ^19^ ^20,21^. Although LHb neuronal mechanisms have been extensively studied^22^, how astrocytes respond to early-life stress and how they influence local synaptic and neuronal function, remain largely unexplored.

Astrocytes regulate neuronal function through multiple mechanisms, among which connexins (Cx) based glial networks stand out as a critical mediator of intercellular communication and circuit homeostasis. There are 21 members of connexins in humans (20 in mice), of which Cx43 and Cx30 are predominant in astrocytes ^23–25^, primarily forming gap junction channels that integrate macroglial cells into panglial syncytia. Gap junctions facilitate the exchange of ions, metabolites, and signalling molecules and contribute to metabolic and functional support of neurons. Connexins can also form hemichannels, which are activated in pathological conditions and may mediate aberrant gliotransmission ^26–29^. Specifically, in models of stress-induced depression, astrocytic connexins expression and function are altered in a region-specific manner. In the ventral hippocampus, astrocytic Cx43 hemichannels are activated, leading to increased extracellular glutamate in stress-induced depression ^30^, whereas in the prefrontal cortex, gap junction coupling is reduced ^31,32^. In the early life stress due to parental separation, expression of Cx43 and Cx30 in hippocampal astrocytes is decreased, along with augmented Ca^2+^ signalling and neuronal activity ^33^. Despite these region-specific observations, a critical developmental gap remains whether connexins in the LHb astrocytes are disrupted in response to early-life stress, and how such disruption contributes to depressive pathology remains entirely unknown.

To address the above knowledge gap, we employed our well-characterised mouse model of early life parental separation ^33^. In this model we found an LHb astrocytic pathological cascade in which reduced Cx43 level and compromised gap junctional coupling promote astrocyte atrophy. Single-cell sequencing revealed distinct transcriptional states of LHb astrocytes following parental separation. Notably, cell-cell communication analysis further predicted reduced astrocyte-neuron communication and remodelling of adhesion-associated signalling pathway. Indeed, by super-resolution imaging we observed aberrant synaptic structure, which translated into reduced synaptic transmission that ultimately drives depressive-like behaviours. Notably, despite the reduction in both excitatory and inhibitory synaptic inputs, LHb neurons exhibited elevated Ca^2+^ dynamics, suggesting pathologically increased intrinsic neuronal excitability. Mechanistically, stress-induced Cx43 dysregulation affects astrocyte structure through Ca^2+^-dependent Rac1 GTPase activation and reduced levels of phosphorylated Ezrin (p-Ezrin). Our findings reveal an LHb astrocytic Cx43 dependent mechanism triggered by early life stress; this mechanism couples astrocyte structure to synaptic organization and behavioural vulnerability, redefining adolescent depression as a disorder driven in part by glial network failure rather than neuronal malfunction alone.

## Results

### LHb astrocytes exhibit a specialised regional phenotype

To determine whether LHb astrocytes possess region-specific features, we first compared the morphology of astrocytes in LHb with other brain regions implicated in mental disorders, including the amygdala, hippocampus, prefrontal cortex (PFC) and lateral hypothalamus (LH) from wild-type (WT) mice by measuring their territory, circularity (Circ.), aspect ratio (AR) and solidity. We then assessed the correlation coefficient for the morphological z score, which standardises each morphological parameter across all astrocytes from all brain regions, and applied principal component analysis. LHb astrocytes emerged as a morphologically distinct population, with a z score of -1.29 in area and 0.56 in solidity, distinguishing them from astrocytes in other brain regions by their compact territories and increased solidity (Fig 1a).

**Fig 1.**
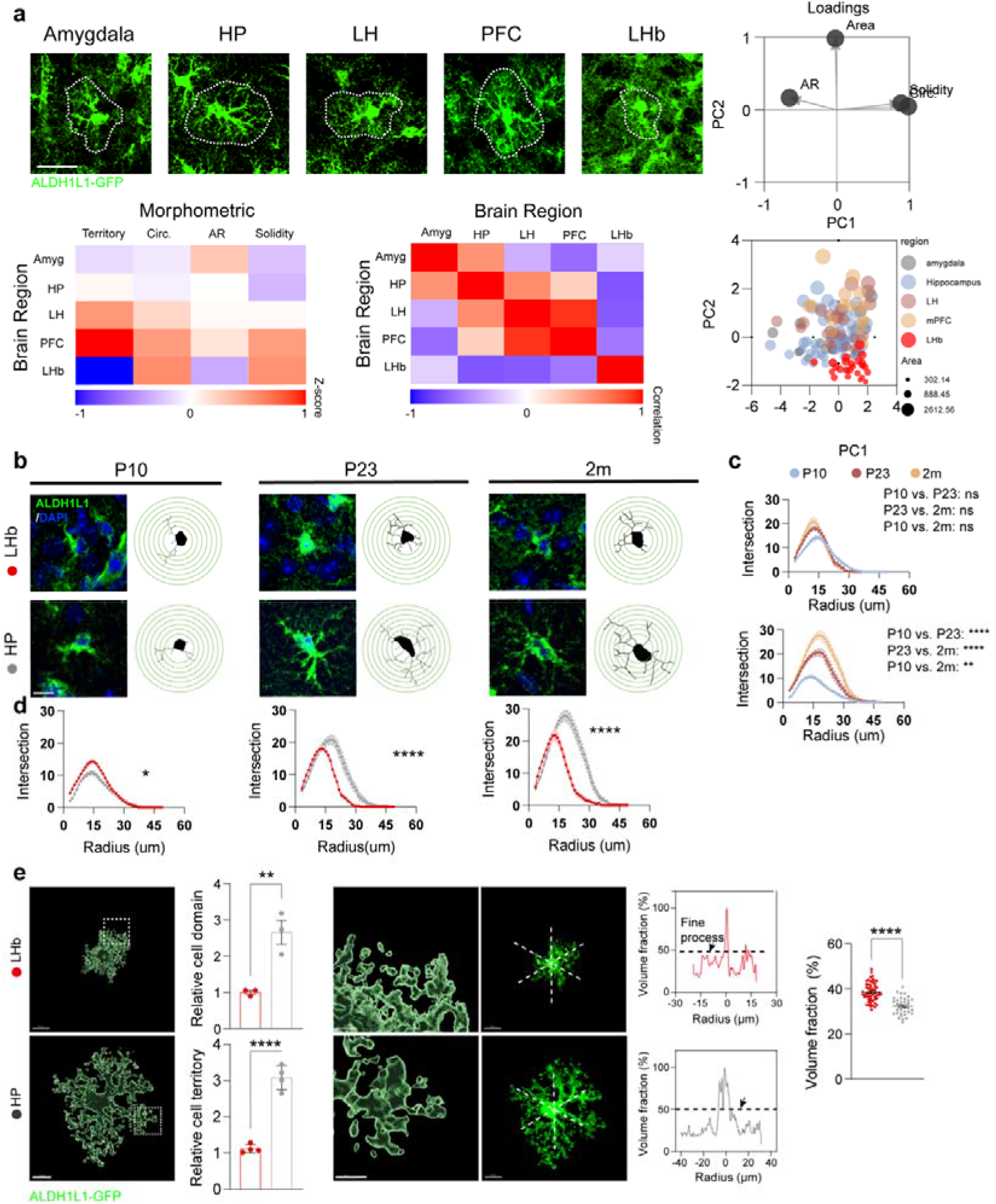
Astrocytes in LHb exhibit distinct morphologic features. (a) Morphological analysis of astrocytes across amygdala, HP, LH, PFC, and LHb. Top left: Fluorescence image of ALDH1L1-GFP. Top right: PCA loading plot illustrating the contribution of morphometric parameters including area, complexity, AR, and circularity. Bottom left: Heatmap depicting Z-scored morpho9metric features across brain regions. Bottom right: Correlation matrix of morphological features between brain regions (n = 3 mice, Scale bar = 20 μm). (b) Immunofluorescence images of ALDH1L1 labelled astrocytes and their Sholl features in HP and LHb (n = 15-35 cells from 3-4 mice, Scale bar = 10 μm). (c) The intersection number analysis from Sholl in each region across different ages (P10, P23, and 2 months). (d) The intersection number assessed from Sholl analysis in each age and the comparison between HP and LHb. (e) Morphology of astrocytes in LHb and HP: 3D reconstruction of ALDH1L1-labeled astrocytes fluorescence images, the quantification of its cell domain (n = 3 mice, Scale bar = 10 μm), the 3D images of leaflet details, the fluorescence images with crossed line to illustrate volume fraction analysis, and the quantification of volume fraction between HP and LHb (LHb, n = 56 cells from 3 mice, HP, n = 34 cells from 3 mice, Scale bar = 5 μm).

To visualise LHb astrocytes, we immunostained astrocytes with antibodies against brain lipid-binding protein (BLBP), glial fibrillary acidic protein (GFAP), and aldehyde dehydrogenase 1 family member L1 (ALDH1L1) at postnatal day 10 (P0), P23, and 2 months of age, representing childhood, adolescence, and adulthood. Immunoreactivity of ALDH1L1 in LHb maintained at a relatively high level across all ages, BLBP staining significantly declined during postnatal development, whereas GFAP immunoreactivity was low at all ages (Supplementary Fig. 1a). Thus, we used ALDH1L1 immunolabelling and ALDH1L1-GFP reporter mice for analysing LHb astrocytes in the subsequent experiments.

Astrocytic morphological diversity is closely linked to their function. To assess the complexity of astrocytic branches (the primary arborisation ^34^), we performed Sholl analysis of ALDH1L1-positive astrocytes at different ages in the LHb and hippocampus (both associated with neuropsychiatric diseases ^35^). The complexity of astrocytes in the hippocampus increased with age, whereas in the LHb, astrocytic complexity remained unchanged (Fig. 1b, c). At P10, astrocytes in the hippocampus exhibited lower complexity compared to those in the LHb. However, at P23 and 2 months, astrocytic branches in the LHb were significantly less complex compared to the hippocampus (Fig. 1b, d). When comparing astrocytes in the LHb with those in the hippocampus at P23, the radius at max intersection was significantly shorter, and the number of total intersections was significantly lower (Extended Data Fig. 1b). To quantify the size of astrocytes, we measured their territories (z-stack maximal projection images) and astrocytic domains (as 3D reconstruction of z-stack images). Notably, although both territory and domain of astrocytes in LHb were significantly smaller than in hippocampus (Fig. 1e), the volume fraction of leaflets^36^, estimated by measuring fluorescence in ALDH1L1-GFP reporter mice, was significantly higher in LHb (38.45% ± 0.56%, n = 56 cells from 3 mice) than that in hippocampus (32.31% ± 0.67%, n = 34 cells from 3 mice; P < 0.0001, Fig. 1e). Thus, LHb astrocytes, when compared to astrocytes from other brain regions, have more compact morphology but more developed leaflets.

Connexins staining was quantified by measuring the fluorescence intensity of immunolabelled Cx30 and Cx43 within ALDH1L1-GFP^+^ astrocyte. Levels of both Cx43 and Cx30 in the LHb astrocytes were significantly higher than in the hippocampus (Fig. 2a, b). Accordingly, the dye coupling among LHb astrocytes was greater than that in the hippocampus (HP: 3.17 ± 0.60, n = 3, LHb: 15.83 ± 1.89, n = 3; P < 0.01) (Fig. 2c). The ethidium bromide (EtBr) uptake in both LHb and hippocampal astrocytes was very low (Fig. 2d), suggesting that hemichannels in both regions were not active under physiological conditions.

**Fig 2.**
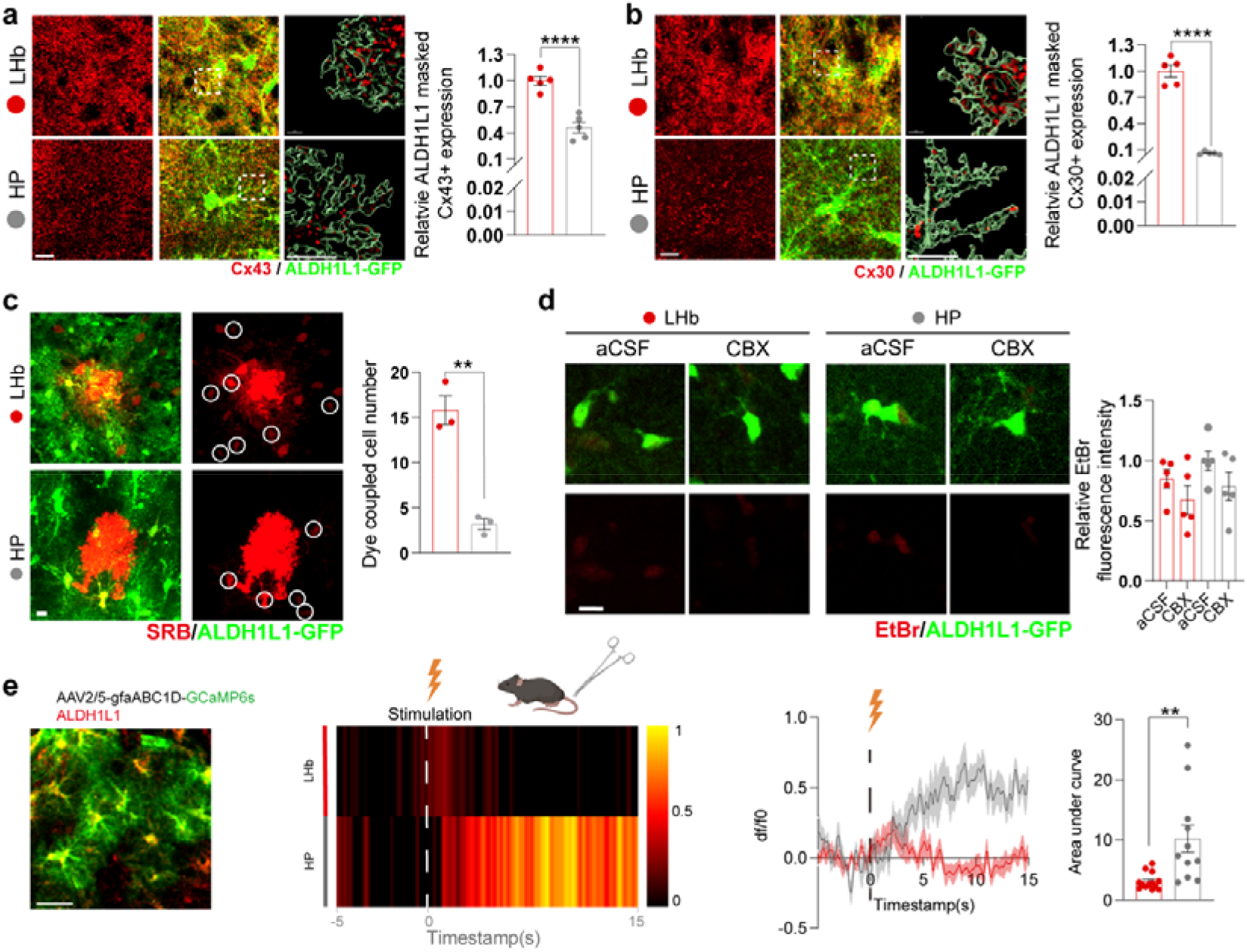
Higher expression of Cx and strengthen gap junction coupling in LHb astrocytes. (a) Representative image of Cx43 staining and ALDH1L1-GFP, 3D reconstruction, and quantification in the LHb and HP (n = 5 mice, Scale bar, 10 μm). (b) Representative images and quantification of Cx30 staining and ALDH1L1-GFP, 3D reconstruction and quantification in the LHb (n = 5 mice, Scale bar, 10 μm). (c) Representative images and quantification of dye diffusion through patch clamp injection (LHb, n = 3 mice, HP, n = 3 mice, Scale bar, 10 μm). (d) Representative image of EtBr and ALDH1L1-GFP staining and quantification of EtBr uptake by ALDH1L1-GFP labelled astrocytes (n = 5 mice, Scale bar, 10 μm). (e) *In vivo* fiber photometry recording of astrocytic calcium response to tail clamping: Representative fluorescence images of GCaMP6s and ALDH1L1 immunostaining, the heatmap of average dF/F_0_ during recordings and its curve, and the quantification of area under curve (LHb, n = 10 stimulations from 3 mice, HP, n = 11 stimulations from 3 mice, Scale bar, 10 μm).

To further characterise the functional dynamics of LHb astrocytes, we performed *in vivo* fibre photometry to monitor astrocytic Ca^2+^ responses to tail clamping. Notably, LHb astrocytes exhibited a significantly shorter Ca^2+^ rise time compared to hippocampal astrocytes, indicating a faster responsiveness to incoming signals. However, the peak amplitude of the Ca^2+^ transient was significantly lower in LHb astrocytes than in hippocampal ones (Fig 2e).

In summary, LHb astrocytes show prominent regional specialisation: they are smaller in overall size with fewer primary branches but denser fine processes; they have abundant Cx43 and Cx30 levels supporting strong gap-junction coupling, and faster Ca^2+^ response to stimulation. These features suggest that LHb astrocytes can form a compact yet efficient tripartite synapse architecture. The unique morphology and molecular profile of LHb astrocytes imply that they may be selectively vulnerable to stress. This raises the question of how pathological conditions, particularly early-life stress, might disrupt this specialised astrocytic network, for example by targeting Cx43-mediated connectivity.

### Morphological and functional remodelling of LHb astrocytes in response to early **life stress**

To determine whether LHb astrocytes are susceptible to early□life stress, we used parental separation to model early-life stress in mice ^33^, and assessed their behavioural and astrocytic phenotype at P23. In the Open Field Test (OFT) for anxiety-like behaviours, mice in the Separation group were less active in the central area, while the total distance travelled was similar to the Control mice (Fig. 3a). In the Forced Swim Test (FST) which assesses depressive-like behaviours, mice in the Separation group spent more time swimming but less time struggling (Fig. 3b), indicating a predominant immobile state.

**Fig 3.**
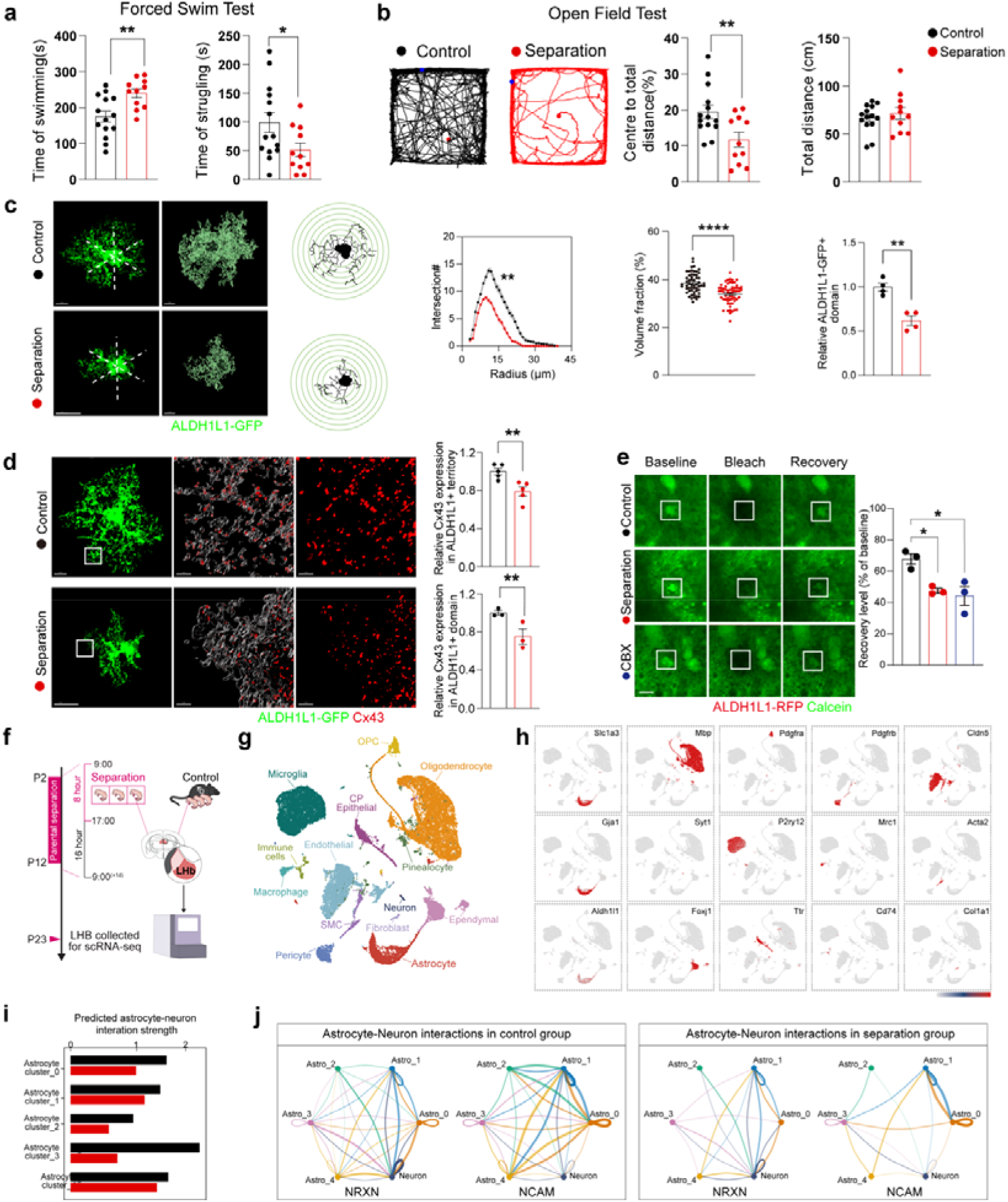
Morphology, molecular and functional changes of astrocytes in response to parental separation. (a) Mice behaviour in forced swim test (Control n = 14 mice, Separation n = 11 mice). (b) Mice behaviour in open field test (Control n = 14 mice, Separation n = 11 mice). (c) Morphological changes of astrocytes in LHb: Representative fluorescence images of LHb astrocytes labeled by ALDH1L1-GFP and its 3D reconstruction, the skeletonised images and sholl features, profiles of ALDH1L1-GFP fluorescence in volume fraction analysis and the quantification of volume fraction (Control, n = 56 cells from 3 mice, Separation, n = 61 cells from 3 mice), cell domain (n = 4 mice) and Sholl analysis (Control n = 45 cells from 3 mice, Separation n = 49 cells from 3 mice, cell domain: n = 4 mice, Scale bar, 10 μm). (d) Representative images of Cx43, ALDH1L1-GFP co-staining in the LHb, and quantification of Cx43 level in ALDH1L1 positive territory and domain (territory n = 5 mice, domain n = 3 mice, Scale bar, 10 μm). (e) Representative image of Calcein staining during gap-FRAP assay (n = 3 mice, Scale bar, 10 μm). (f) Experimental design. Mice underwent maternal separation during the early postnatal period, and LHb tissue was collected at P23 for single-cell RNA sequencing. Each condition was generated from pooled tissue from multiple animals. (g) UMAP representation of major cell populations identified in the LHb dataset, including astrocytes, oligodendrocytes, oligodendrocyte precursor cells (OPCs), microglia, macrophages, immune cells, neurons, endothelial cells, pericytes, smooth muscle cells (SMCs), fibroblasts, ependymal cells, choroid plexus (CP) epithelial cells and pinealocytes. (h) Heat map of relative pathway activity across astrocyte subtypes in control and separation conditions. Representative pathways are grouped into metabolic/stress, inflammatory and survival-associated programs. (i) CellChat analysis of predicted astrocyte–neuron communication. (j) Circle plots showing NRXN- and NCAM-associated communication networks. Nodes represent astrocyte subtypes and neurons, and edge width represents inferred communication strength.

The complexity of primary arborisation of ALDH1L1-GFP astrocytes in the LHb was significantly decreased in the Separation group, with astrocytic domains decreased by 38.07% ± 7.03% (n = 4; P < 0.01) compared to the Control group (Fig. 3c). In addition, we found a significant decrease in leaflet volume fraction in the Separation group by 4.47% ± 0.78% (Control, n = 56 cells from 3 mice, Separation, n = 61 cells from 3 mice; P < 0.0001) (Fig. 3c). To assess whether this astrocytic remodelling was cell□type specific, we also examined other glial cell types, including microglia, oligodendrocytes, and oligodendrocyte precursor cells, and found no overt morphological changes (Extended Data Fig. 2a), confirming that astrocytes are the primary glial responders to stress□induced changes in the LHb.

The presence of Cx43 and Cx30 in astrocytes was assessed in ALDH1L1-GFP+ astrocytic territory and domain, followed by a 3D reconstruction of connexin distribution within an individual astrocyte. The Cx43 levels were significantly reduced in mice that underwent parental separation, both within ALDH1L1-GFP+ territory (decreased by 21.05% ± 5.78%, n = 5; P < 0.01) and domain (decreased by 38.94% ± 10.79%, n = 3; P < 0.05) (Fig. 3d). This downregulation of Cx43 did not appear until P23 (Extended data Fig 2b). Conversely, Cx30 level remained unaffected (Extended Data Fig. 2c). To determine whether selective Cx43 loss results in functional deficits, we perform fluorescent recovery after photobleaching (gap-FRAP) to quantify gap junctional communication, and found that fluorescence recovery was significantly reduced in astrocytes in the Separation group (Control vs. Separation: 63.44% ± 3.34% vs. 47.46% ± 1.50%, n = 3; P < 0.01), suggesting decreased gap junctional coupling (Fig. 3e). This was further corroborated by a reduced dye coupling in the dye injection experiments (Control vs. Separation: 15.20 ± 1.66 vs. 4.429 ± 0.90, Control n = 5 cells from 3 mice, Separation n = 7 cells from 3 mice, P < 0.001) (Extended Data Fig. 2d). The EtBr uptake by astrocytes in the LHb was not altered in the Separation group, suggesting that hemichannels in LHb astrocytes remained inactive (Extended Data Fig. 2e).

To define cellular changes in the LHb following early-life stress, we performed single-cell RNA sequencing of LHb tissue from control and maternal separation mice (Fig. 3f). Major neuronal, glial, vascular, and immune populations were resolved by unsupervised clustering and annotated using canonical markers (Fig. 3g, h; Extended Data Fig. 2f, g). Focus on astrocytes revealed marked transcriptional differences between control and separation conditions (Fig. 3i, j). Astrocyte-specific clustering identified five transcriptional states (Fig. 3k). Their relative representation differed between the control and separation datasets. Pathway analysis further revealed state-dependent changes across metabolic and stress-related, inflammatory, and survival-associated transcriptional programs (Fig. 3l). These data indicate that maternal separation is associated with a substantial reorganization of the astrocyte transcriptional landscape in the LHb.

Astrocyte–neuron interactions at tripartite synapses are shaped in part by cell-adhesion molecules ^37^. We therefore examined whether astrocyte-state remodelling after maternal separation was accompanied by altered intercellular communication. CellChat predicted reduced astrocyte-to-neuron communication across all astrocyte states (Fig. 3m). Among adhesion-associated pathways, NRXN and NCAM showed prominent network remodelling. Both networks were broadly reduced after separation, with marked losses of astrocyte–astrocyte and astrocyte-to-neuron communication (Fig. 3n). NRXN-associated communication also showed reduced neuron–neuron connectivity, whereas neuron-to-astrocyte signalling was relatively preserved. Together, these changes suggest that maternal separation is affects adhesion-related communication at the astrocyte–neuron interface, indicating altered molecular organization of tripartite synapses.

Taken together, early-life parental separation-induced depressive behaviour in mice was associated with structural atrophy and reduced complexity of astrocytes, as well Cx43 downregulation, resulting in decreased gap junctional communication. Thus, early-life stress selectively compromises Cx43-dependent LHb astrocytic networks, providing an upstream glial basis for subsequent synaptic and neuronal malfunction.

### Impaired tripartite synapse structure and function in the LHb in mice with parental separation

Astrocytes extend leaflets to closely interact with synapses in spatial proximity, contributing to a tri-(multi)-partite synapse and astroglial cradle through which astrocytes support and regulate synaptic transmission ^38–40^. Astrocyte atrophy and reduced leaflets decrease their association with synapses, resulting in synaptic malfunction ^41^. Synapses in the LHb are predominantly excitatory ^42,43^, although inhibitory synapses also exist ^44^. Our analysis revealed that 83.57% ± 2.82% (n = 3 mice) of synapses in LHb were Synapsin and Homer1 positive, suggestive of excitatory synapses, while only 16.43% ± 2.82% (n = 3 mice) were inhibitory as judged by Synapsin and Gephyrin co-staining (Extended Data Fig. 3). We therefore systematically examined the association of astrocytes with both excitatory and inhibitory synapses.

First, we investigated the association of astrocytes with excitatory synaptic sites. VGLUT2 and Homer1 were used to stain the pre- and post-excitatory synaptic compartments, respectively, while ALDH1L1-RFP fluorescence reporter mice were used to visualise astrocytes. Images were analysed with super-resolution microscopy, followed by 3D reconstruction (Fig. 4a). Triple co-localisation of ALDH1L1-RFP, Homer1, and VGLUT2 was reduced by 52.9% ± 11.81% (n = 5, P < 0.01) in mice that experienced early-life parental separation (Fig. 4a). Numbers of both VGLUT2 and Homer1 puncta within an individual astrocyte domain were also significantly decreased (VGLUT2 by 38.96% ± 7.35%, P < 0.001, Homer1 by 46.85% ± 8.72%, n = 5, P < 0.001) in mice with parental separation (Extended Data Fig. 4a). Thus, the association between astrocytes leaflets and excitatory synapses was reduced at both pre- and post-synaptic sites. However, the total number of VGLUT2 and Homer1 co-localised puncta was not affected by parental separation (Extended Data Fig. 4a), suggesting preserved synaptic density.

**Fig 4.**
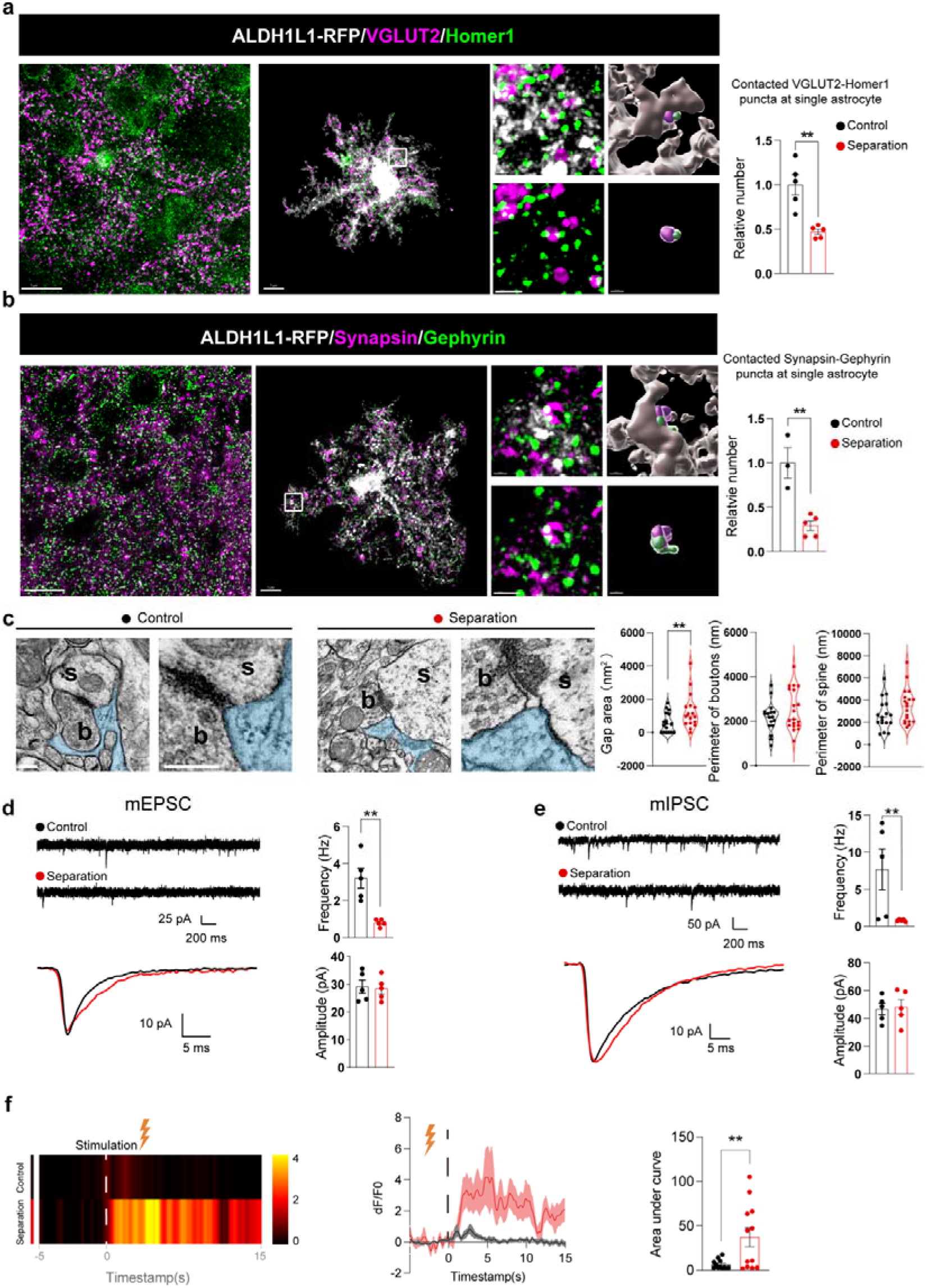
Parental separation disrupts synaptic structure and function and alters neuronal Ca^2+^ activity. (a) Representative images of ALDH1L1-RFP, VGLUT2 and Homer1, and the number of RFP-labelled astrocytes co-stained with VGLUT2 and Homer 1 in LHb (Control n = 5 mice, Separation n = 5 mice, Scale bar, left, 10 μm, right, 1 μm). (b) Representative images of ALDH1L1-RFP labelled astrocytes co-stained with Synapsin, and Gephyrin, and the number of co-localisation puncta of these markers in LHb (Control n = 3 mice, Separation n = 5 mice, Scale bar, left, 10 μm, right, 1 μm). (c) Representative electron microscope images of multipartite synapse and the gap area between astrocytes leaflets and synaptic cleft, the perimeter of boutons, and spines in Control and Separation groups (Control n = 17 cells from 4 mice, Separation n = 17 cells from 4 mice, Scale bar, 200 nm). (d) Representative current traces pattern and average mEPSC and quantification of amplitudes and frequencies (n = 5 cells from 3 mice). (e) Representative current traces pattern and average mIPSC (e, n = 5 cells from 3 mice) and quantification of amplitudes and frequencies (f) Neuronal Ca^2+^ dynamics in parental separation model LHb: The heatmap of average df/f0 during recordings and its curve, and the quantification of area under curve (Control n = 12 stimulation from 5 mice, Separation n = 12 stimulation from 4 mice).

Similar analysis was performed for inhibitory synapses (Fig. 4b). The co-localisation of ALDH1L1-RFP, Synapsin (inhibitory pre-synaptic marker) and Gephyrin (inhibitory post-synaptic marker) was significantly decreased by 70.68% ± 14.45% (Control n = 3, Separation, n = 5, P < 0.01) in mice with parental separation (Fig. 4b). Both Synapsin and Gephyrin puncta within a single astrocyte were decreased (Extended Data Fig. 4b, Synapsin: decreased by 59.26% ± 8.87%, P < 0.001, Gephyrin: decreased by 73.64% ± 11.26%, Control n = 3, Separation, n = 5, P < 0.001) in mice with early life stress, suggesting decreased association of astrocytes leaflets with both inhibitory pre- and post-synaptic compartments. The total number of Synapsin and Gephryin co-localised puncta was not affected (Extended Data Fig. 4b). Thus, early-life parental separation stress impairs the relations between astrocytes and synapses in the LHb, without affecting the density of synaptic contacts.

Spatial correlation of astrocytes and synapses was further investigated through transmission electron microscopy (TEM) ^45–48^. Gap area between astrocytes leaflet and neuronal synapses, was significantly increased by 679.8 ± 293.2 nm^2^ (n = 17 cells from 4 mice, P < 0.05) in mice that underwent parental separation. Perimeters of boutons and spines were not affected. These data highlight aberrant tripartite synapse structure in the LHb in the Separation group (Fig. 4c). Parental separation therefore selectively disrupts astrocytic leaflets engagement with synaptic compartments, despite preservation of overall synapses numbers.

We next investigated whether structural disruption of tripartite synapses translates into functional synaptic deficits. Whole-cell patch-clamp recordings from LHb neurons were performed to assess synaptic transmission. The frequency of mEPSC was significantly decreased in neurons from the Separation group, although the mEPSC amplitude did not differ from Control group, which was further reflected by the averaging of recorded mEPSC events (Fig. 4d). Similar decrease in frequency (but not in amplitude) of mIPSC was observed in the Separation group (Fig. 4e), suggesting an impaired synaptic function implying a deficient perisynaptic neurotransmitter release in the LHb in response to early-life stress.

To determine how this synaptic weakening affects neuronal function, we performed *in vivo* photometry to record LHb neuronal Ca^2+^ activity. Surprisingly, despite the reduced synaptic inputs, LHb neurons in the Separation group showed significantly increased Ca^2+^ transients compared to Controls (Fig. 4f), indicating that neuronal activity was enhanced. This suggests that the primary driver of elevated LHb neuronal activity in adolescent depression is not an increased synaptic input, but rather an aberrant increase in intrinsic neuronal excitability, potentially due to compromised astrocytic support following reduction of Cx43 gap junction astrocytic syncytial connectivity.

In summary, these findings demonstrate that early-life parental separation disrupts the structural and functional integrity of synapses in the LHb, reduces both excitatory and inhibitory synaptic transmission, but elevates neuronal Ca^2+^ activity. This uncoupling between synaptic input and neuronal output establishes a pathological state in which LHb neurons become hyperactive despite weakened synaptic drive, setting the stage for the depressive□and anxiety□like behaviours observed in these mice, and raising the question of whether astrocytic Cx43 loss is sufficient to recapitulate this entire phenotype.

### Astrocytic Cx43 loss is sufficient to recapitulate early-life stress induced pathology

To determine whether reduced astrocytic Cx43 and gap junctions malfunction are causally responsible for, rather than merely associated with, the pathology observed in the aforementioned mice with early life parental separation, we injected AAV2/5-ALDH1L1-shRNA-Gja1-RFP into the LHb to selectively knock down Cx43 in astrocytes, with RFP fluorescence reporting ALDH1L1-positive astrocytes. Scrambled shRNA was used as a control (Fig. 5a). The specificity of virus infection in LHb astrocytes and efficacy of astrocytic Cx43 knockdown were confirmed in untreated naïve mice using immunostaining, which demonstrated that 99.27% ± 0.46% and 99.44% ± 0.23% of RFP-positive cells were astrocytes (Extended Data Fig. 5a). Cx43 staining was decreased by 97.63% ± 6.1% in Cx43 knockdown mice (n = 4, P < 0.0001, Extended Data Fig. 5b).

**Fig 5.**
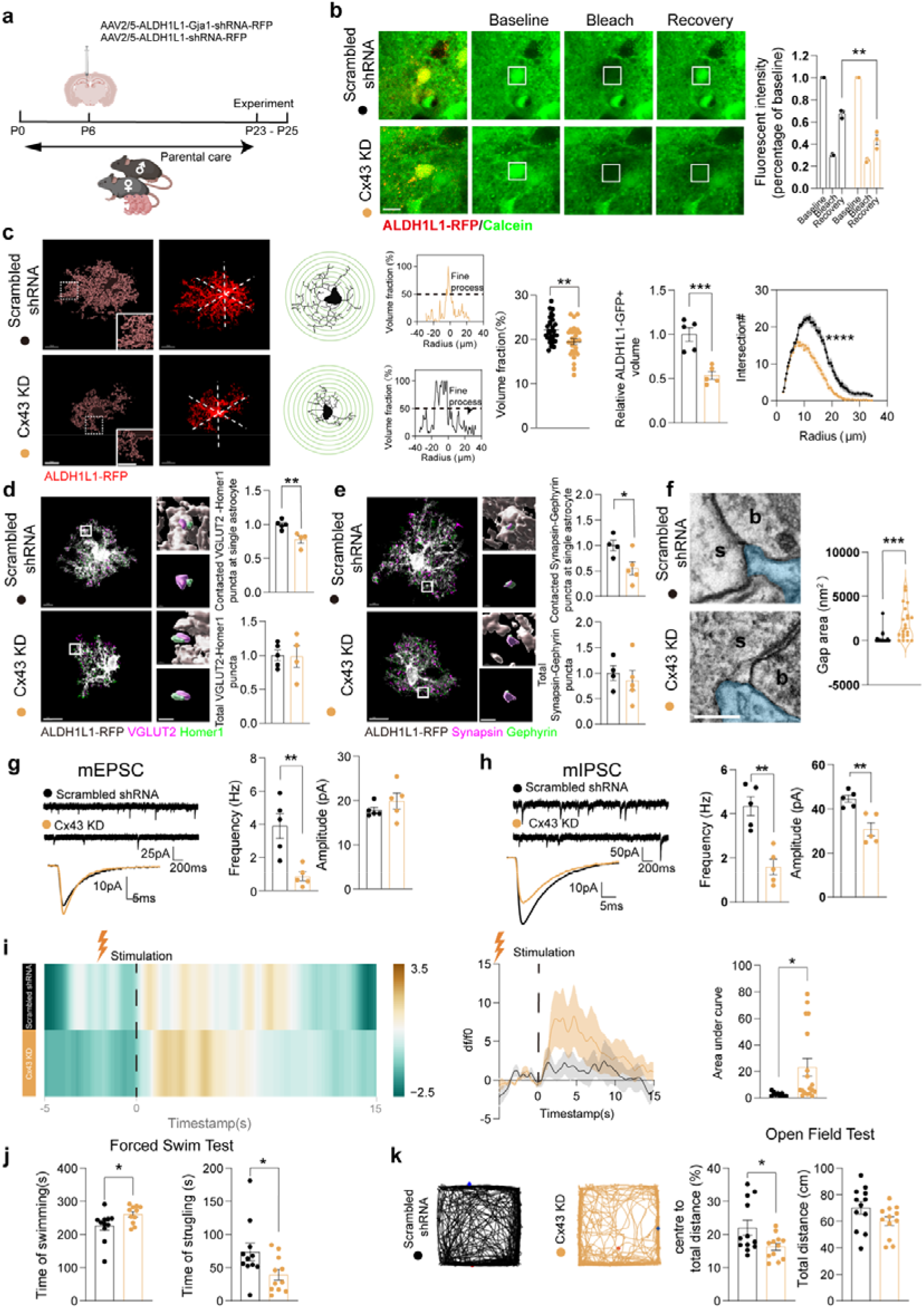
Cx43 downregulation in the LHb directly drives the phenotype of parental separation model. (a) Schematic paradigm of Cx43 knockdown in wildtype mice. (b) Representative images and quantification of Calcein during fluorescence recovery after photobleaching assay (n = 3 mice). (c) Morphologic features of astrocytes: Representative fluorescence images of LHb astrocytes labelled by ALDH1L1-RFP and its 3D reconstruction, the morphology traces and Sholl features, profiles of ALDH1L1-RFP fluorescence in volume fraction analysis and the quantification of volume fraction (Scrambled shRNA, n = 29 cells from 3 mice, Cx43 KD, n = 27 cells from 3 mice), cell domain (n = 5 mice) and Sholl analysis (Scrambled shRNA n = 34 cells from 5 mice, Cx43 KD n = 33 cells from 4 mice, Scale bar, 10 μm). (d) Representative images of ALDH1L1-RFP labelled astrocytes co-stained with VGLUT2 and Homer1, and the number of triple co-stained puncta, as well as VGLUT2 and Homer1 co-localisation in LHb (Scrambled shRNA, n = 5 mice, Cx43 KD, n = 4 mice, Scale bar, left, 10 μm, right, 1 μm). (e) Representative images of ALDH1L1-RFP labelled astrocytes co-stained with Synapsin, and Gephyrin, and the number of triple co-localisation puncta of these markers, as well as Synapsin and Gephyrin co-localisation (Scrambled shRNA, n = 4 mice, Cx43 KD, n = 5 mice, Scale bar, 10 μm, right, 1 μm). (f) Representative electron microscope images of multipartite synapse and quantification of the gap area between astrocytes and synaptic cleft (Scrambled shRNA n = 21 cells from 3 mice, Cx43 KD n = 21 cells from 4 mice, Scale bar, 200 nm). (g) Representative current traces and average curve of mEPSC and the quantification of amplitude and frequency (n = 5 cells from 3 mice). (h) Representative current traces and average curve of mIPSC and the quantification of amplitude and frequency (n = 5 cells from 3 mice). (i) Heatmap and averaged F/F traces of LHb astrocyte Ca^2+^ responses to stimulation, with quantification of the area under the curve (AUC) during *in vivo* fibre photometry recordings (Scrambled shRNA n = 9 stimulation form 3 mice, Cx43 KD, n =18 stimulation from 6 mice). (j) Forced swim test of mice with or without astrocytic Cx43 knockdown in LHb (n = 11 mice). (k) Representative traces of mice in Open field test, and quantification of percentage of mice travelling in centre area, as well as the total distance (n = 12 mice).

We first asked whether astrocytic coupling was disrupted following Cx43 knockdown. Gap-FRAP analysis revealed a significant reduction in Calcein fluorescence recovery level (Scrambled shRNA vs. Cx43 knockdown: 77.60% ± 5.95% vs. 48.65% ± 6.50%, n = 3, P < 0.01) following Cx43 knockdown (Fig. 5b), indicating a suppression of astrocytic gap junctional coupling in LHb, which closely resembled the alterations observed in mice subjected to parental separation.

The Cx43 knockdown significantly reduced LHb astrocytes complexity and territorial domain (Fig. 5c). The volume fraction of LHb astrocytes leaflets was also significantly decreased (Scrambled shRNA vs. Cx43 knockdown, 22.19% ± 0.53% vs. 19.48% ± 0.68%, Scrambled shRNA, n = 29 cells from 3 mice, Cx43 knockdown, n = 27 cells from 3 mice, P < 0.01) in Cx43 knockdown mice (Fig. 5c). Thus astrocyte-specific Cx43 knockout recapitulated the phenotype of astrocytes observed following parental separation, and further supported the notion that Cx43 is required for maintaining the highly specialised morphology of LHb astrocytes.

Given the close structural association between astrocytes and synapses, we next examined whether Cx43 downregulation compromises tripartite synapse organisation. In Cx43 knockdown mice, we found a significant decrease in co-localisation of ALDH1L1-RFP, VGLUT2, and Homer1 (by 21.89% ± 5.64%, Scrambled shRNA, n = 5, Cx43 knockdown, n = 4, P < 0.01) in the LHb (Fig. 5d, Extended Fig 5c). Co-localised VGLUT2 or Homer1 puncta at an individual astrocyte was also reduced (VGLUT2: decreased by 28.80% ± 11.45%, P < 0.05, Homer1: decreased by 37.86% ± 12.43%, Scrambled shRNA, n = 5, Cx43 knockdown, n = 4, P < 0.05), while the number of VGLUT2 and Homer1 co-localisations was not changed, indicating that total number of excitatory synapses remained unaffected (Fig. 5d). Similarly, the associations between ALDH1L1-RFP positive astrocytes, Synapsin and Gephyrin in the LHb were significantly decreased (by 44.83% ± 17.28%, Scrambled shRNA, n = 4, Cx43 knockdown, n = 5, P < 0.05) in Cx43 knockdown mice, and number of Synapsin (but not Gephyrin) that contacted at a single astrocyte were also decreased by 58.03% ± 10.34% (Scrambled shRNA. n = 4, Cx43 knockdown, n = 5, P < 0.001) (Fig. 5e). However, no changes were observed in the total Synapsin and Gephyrin co-localisation, suggesting preserved number of inhibitory synapses (Fig. 5e, Extended Fig. 5d). There was a significant increase in the gap area between astrocytes and synapses (by 1,748 ± 440.5 nm^2^, Scrambled shRNA n = 21 cells from 3 mice, Cx43 knockdown n = 21 cells from 4 mice, P < 0.001) and unchanged boutons and spine perimeter in Cx43 knockdown mice under TEM, indicating a disconnection of astrocytes and synapses (Fig. 5f, Extended Fig. 5e).

We next examined whether these structural alterations translated into functional defects. Whole-cell patch-clamp showed that the frequency of mEPSC decreased while the amplitude was not affected by Cx43 knockdown in the LHb (Fig. 5g). On the other hand, both amplitude and frequency of mIPSC were significantly decreased by Cx43 knockdown in the LHb (Fig. 5h), suggesting impaired synaptic function in the LHb. *In vivo* photometry recordings demonstrated that neuronal Ca^2+^ activity in the LHb was markedly increased following astrocytic Cx43 knockdown (Fig 5i).

Finally, we determined whether selective astrocytic Cx43 deficiency was sufficient to drive behavioural abnormalities. In the FST, swimming time was increased while struggling time was decreased in mice with LHb Cx43 knockdown, indicating depressive-like behaviours (Fig. 5j). In the OFT, mice with LHb Cx43 knockdown had a significant reduction in the percentage of distance travelled in the central area, while the total travel distance was unchanged (Fig. 5k), indicating anxiety-like behaviour.

Collectively, these findings indicate that deficiency of astrocytic Cx43 in the LHb is sufficient to phenocopy the structural, functional, and behavioural abnormalities induced by early-life stress, identifying astrocytic Cx43 as a causal regulator of LHb malfunction.

### Astrocytic Ca^2+^ hyperactivity is required for cytoskeletal remodelling and depressive pathology downstream of Cx43 deficiency

Having established astrocytic Cx43 deficiency as a causal driver of LHb malfunction, we investigated how impaired astrocytic gap junction coupling leads to astrocyte atrophy. One key role of connexin gap junctions in the astroglial network is mediating intercellular Ca^2+^ waves. We therefore examined whether Cx43 deficiency alters astrocytic Ca^2+^ dynamics. *In vivo* photometry revealed that astrocytic Ca^2+^ activity in LHb was significantly increased in both parental separation mice and Cx43 knockdown mice, as evidenced by an increase in area under curve (AUC) (Fig. 6a). Consistently, *ex vivo* slice Ca^2+^ imaging exhibited increased frequency of spontaneous Ca^2+^ transients, greater amplitude of Ca^2+^ events and increased AUC in LHb from Separation group (Extended Data Fig. 6a). Cx43 knockdown group also showed increased Ca^2+^ activity reflected by both *ex vivo* Ca^2+^ imaging and *in vivo* photometry recordings (Extended Data Fig. 6b, c). These findings demonstrate that astrocytic Ca^2+^ hyperactivity represents a common downstream consequence of both parental separation and astrocytic Cx43 deficiency, suggesting that impaired gap junction coupling may be associated with astrocytic pathology induced by early-life stress.

**Fig 6.**
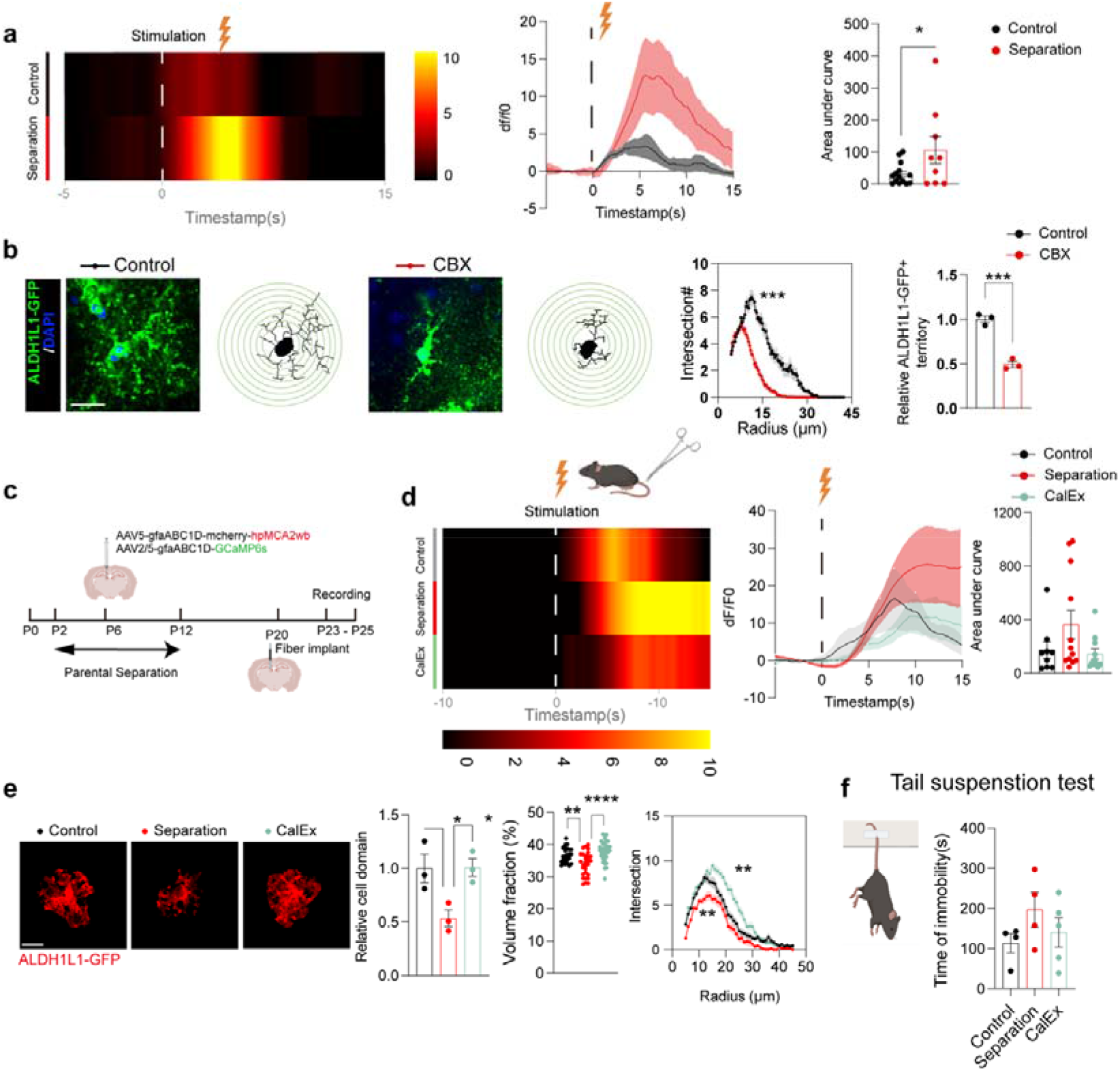
Cx43 gap junction regulates astrocytic morphology through a calcium dependent way. (a) Heatmap and average curve of F/F_0_ signal obtained from photometry recording of astrocytic Ca^2+^ signals in parental separation model LHb, and the quantification of area under curve (Control n = 14 stimulation from 6 mice, Separation, n = 9 stimulation from 3 mice). (b) Representative fluorescence images of ALDH1L1-GFP in slice culture, Sholl analysis and quantification of cell territory (Sholl n = cells from 3 mice, territory n = 3 mice, Scale bar, 20 μm). (c) Schematic illustration of CalEx treatment in parental separation model. (d) Heatmap and curve of F/F_0_ signal obtained from photometry recording of astrocytes Ca^2+^ signals in CalEx treated mice LHb, and the quantification of area under curve (Control n = 12 stimulation from 4 mice, Separation, n = 19 stimulation from 6 mice, CalEx, n = 14 stimulation from 5 mice). (e) Representative fluorescence images of ALDH1L1 immunostaining, and the quantification of its cell territory, volume fraction and Sholl analysis (territory, n = 3 mice, volume fraction, Control n = 26 cells from 3 mice, Separation, n = 29 cells from 3 mice, CalEx, n = 36 cells from 3 mice, Sholl analysis, Control n = 37 cells from 3 mice, Separation, n = 26 cells from 3 mice, CalEx, n = 36 cells from 3 mice, Scale bar, 20 μm). (f) Tail suspension test of parental separation model mice receiving CalEx treatment and the quantification of immobility time (Control, n = 4 mice, Separation, n = 4 mice, CalEx, n = 5 mice, Scale bar, 10 μm).

To confirm the association between Cx43 gap junction function and astrocyte atrophy, we analysed the changes in morphology of astrocytes in acute brain slices with or without blocking gap junctions by carbenoxolone (CBX, a common gap junction blocker). Following 200 μM CBX treatment for 24 hours, the territories of ALDH1L1-GFP astrocytes in the LHb were significantly reduced by 50.65% ± 5.00% (Control: n = 24 cells from 3 mice, CBX n = 33 cells from 3 mice, P < 0.001); similarly, the complexity of major branches of ALDH1L1-GFP astrocytes quantified by Sholl analysis was significantly decreased (Fig. 6b), indicating that disruption of astrocytic gap junction is sufficient to induce astrocytic structural remodelling, medicated by Ca^2+^ signalling.

To clarify how Ca^2+^ signalling affects astrocyte morphology, we measured Rac1, a small GTPase, known to negatively regulate astrocyte morphology ^49^; Rac1 is upregulated by higher intracellular Ca^2+^ ^50,51^. We observed nearly three times increase in Rac1 levels (n = 3, P < 0.05, Extended Data Fig. 6d), in primary astrocytes isolated from the LHb with Cx43 knockdown, suggesting that intracellular Ca^2+^ signalling is positively linked to Rac1 level, which in turn, resulted in astrocytic atrophy.

To confirm the exact mechanism mediating the regulation of astrocyte morphology by Ca^2+^ signalling, 2-APB (a broad Ca^2+^ channel inhibitor ^52^) was applied to acute brains slices to inhibit Ca^2+^ transients (Extended Data Fig. 7a, b). We also found that Rac1-GTP levels were significantly higher in the Separation group. This increase of Rac1-GTP was diminished when Ca^2+^ signals were inhibited by 2-APB (Extended Data Fig. 7c). In addition, the LHb astrocytes domains in both Control and Separation groups were increased after 2-APB treatment (Extended Data Fig. 7d). The complexity of astrocytes was also significantly increased by 2-APB (Extended Data Fig. 7d), with increased leaflet volume fraction (Extended Data Fig. 7d). These findings suggest that excessive Ca^2+^ signalling lies upstream of Rac1 activation and astrocytic atrophy. Rac1 has been shown to regulate p-Ezrin, a plasmalemmal-actin linker that controls astrocytic leaflet architecture ^53^. Knocking down Cx43 significantly reduced the level of p-Ezrin in astrocytes (Extended Data Fig. 7e). Therefore, Cx43 downregulation and gap junction malfunction in the LHb by parental separation are associated with hyperactivation of Ca^2+^ signalling, which in turn increases Rac1 levels and inhibits the downstream p-Ezrin signalling, ultimately resulting in leaflets retraction and astrocyte atrophy. As a result, astrocytic synaptic coverage is reduced, compromising tripartite synapse structure and function translating into behavioural abnormalities.

We next investigated whether astrocytic Ca^2+^ hyperactivity is sufficient to induce structural remodelling *in vivo*, and is required for the development of depressive pathology. We selectively expressed CalEx, a genetically encoded Ca^2+^-extrusion tool that enhances intracellular Ca^2+^ clearance, in the LHb astrocytes to suppress Ca^2+^ signalling *in vivo* ^54^. This strategy enabled us to normalise astrocytic Ca^2+^ signalling and test its causal contribution to the pathological consequences of dysfunctional Cx43 connectivity. *In vivo* photometry recording confirmed that CalEx significantly suppressed astrocytic Ca^2+^ signals in the LHb of mice with parental separation. This astrocytic Ca^2+^ signals suppression effectively reversed astrocytic atrophy, as demonstrated by elevation of cell domain, volume fraction and major branches from Sholl analysis (Fig. 6e). Finally, the inhibition of CalEx on astrocytic Ca^2+^ also reduced the swimming time of in FST in mice with parental separation (Fig. 6f), indicating that the depressive-like behaviour was rescued by Ca^2+^ signal inhibition.

Taken together, these findings demonstrated that impaired astrocytic gap junction coupling induce astrocytic Ca^2+^ hyperactivity which is responsible for early-life parental separation-induced structural remodelling of astrocytes and depressive-like behaviours.

### Restoration of astrocytic Cx43 re-establishes astrocytic homeostasis and reverses early-life stress-induced LHb dysfunction as well as abnormal behaviours

Given that reduced astrocytic Cx43 in the LHb contributes to early-life parental separation associated anxiety and depressive behaviours, we investigated whether restoring astrocytic Cx43 in the LHb can rescue abnormal behaviours. We overexpressed Cx43 in astrocytic specifically in the LHb in the Separation group by injecting AAV2/5-ALDH1L1-P2A-Gja1-RFP virus (as “Cx43 OE” in the Figure), with RFP-only virus as control (as “Separation” in the Figures) (Fig. 7a). Astrocytic Cx43 overexpression in the LHb was confirmed by co-immunostaining with ALDH1L1 and Cx43, where 99.19 ± 0.08 % and 99.16 ± 0.09% of RFP positive cells, respectively, were astrocytes (Extended Data Fig. 8a), while the astrocytic Cx43 level was significantly increased by 29.1% ± 8.77% (n = 5, P < 0.05) compared with mice receiving RFP-only virus (Extended Data Fig. 8b), indicating the high specificity of this virus as well as its efficiency in overexpressing Cx43. Overexpression of Cx43 also significantly increased fluorescence recovery after photobleaching (Separation vs. Cx43 overexpression: 48.46% ± 3.88% vs. 69.69% ± 4.29%, n = 3, P < 0.05, Extended Data Fig. 8c), suggesting increased astrocytic gap junctional connectivity.

**Fig 7.**
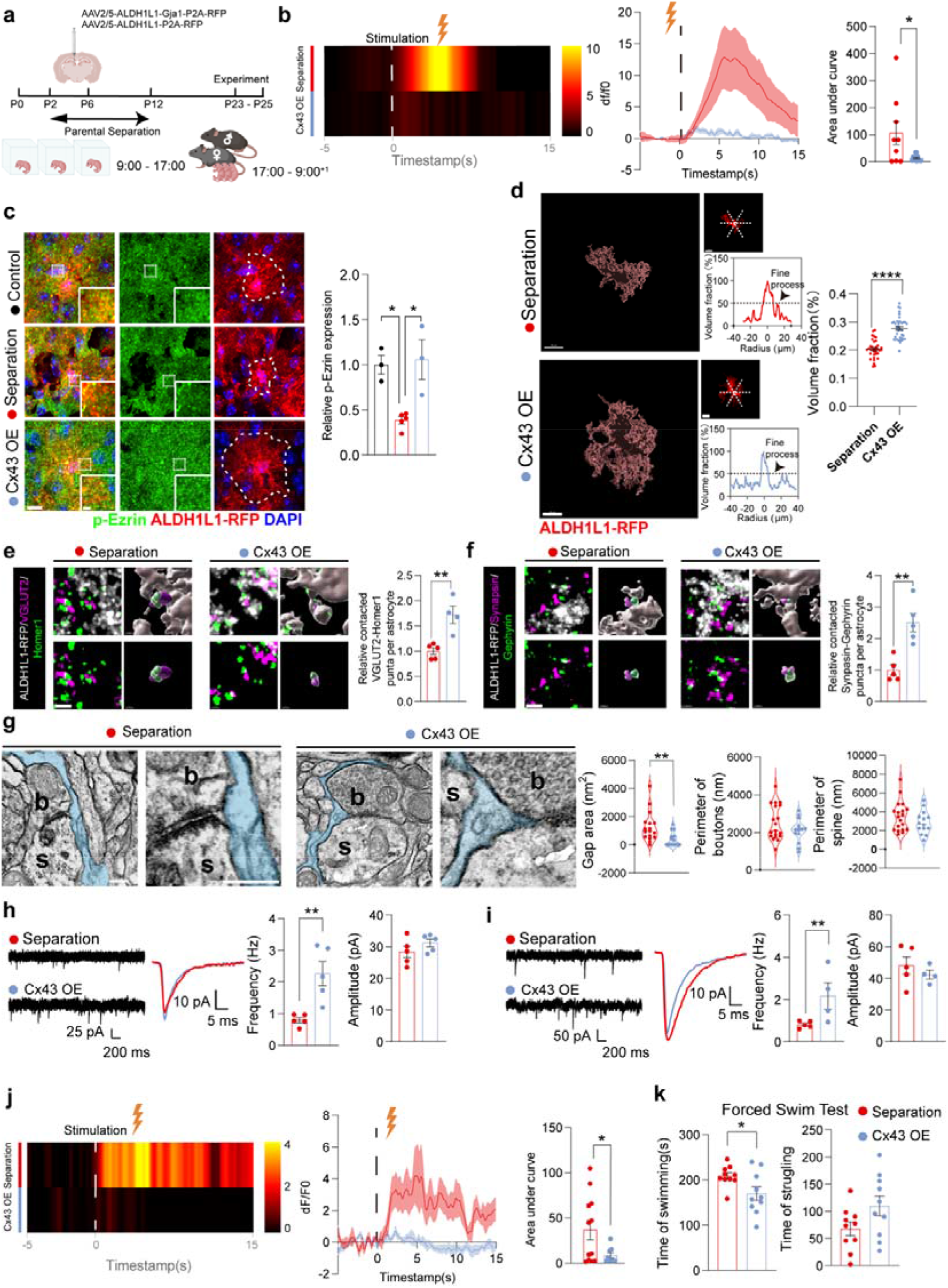
Overexpression of Cx43 in LHb astrocytes alleviates neurological malfunction in Separation model. (a) Experimental paradigm of Cx43 overexpression in Separation model. (b) Heatmap and average curve of astrocytic Ca^2+^ signal in LHb recorded by *in vivo* photometry and the quantification of area under curve (Separation, n = 9 stimulations from 3 mice, Cx43 OE, n = 12 stimulations from 4 mice). (c) Representative images of p-Ezrin and ALDH1L1-RFP and its quantification (Control, n = 3, Separation, n = 5, Cx43 overexpression, n = 3, Scale bar, left, 10 μm, right, 2 μm). (d) Representative images of ALDH1L1-RFP, and the ALDH1L1-RFP fluorescence profiles for volume fraction analysis, and the quantification of leaflet volume fraction in the Separation and Cx43 OE groups (Separation, n = 35 cells from 3 mice; Cx43 OE, n = 39 cells from 3 mice, Scale bar, 10 μm). (e) Representative images, 3D reconstruction and quantification of ALDH1L1-RFP labelled astrocytes co-stained with VGLUT2 and Homer1 in Separation and Cx43 OE mice (Separation n = 5 mice, Cx43 OE, n = 4 mice, Scale bar, 1 μm). (f) Representative images, 3D reconstruction and quantification of ALDH1L1-RFP labelled astrocytes co-stained with Synapsin and Gephyrin (J, Separation n = 5 mice, Cx43 OE, n = 5 mice, Scale bar, 1 μm). (g) Representative electron microscope images and the quantification of gap area between astrocytes and synaptic cleft in LHb (Separation n = 17 cells from 4 mice; Cx43 OE n = 14 cells from 4 mice, Scale bar, 200 nm). (h) Representative mEPSC recordings and averaged mEPSC n Separation and Cx43 OE group mice and its frequency and amplitude (n = 5 cells from 3 mice). (i) Representative mIPSC recordings and averaged mIPSC in Separation and Cx43 OE group mice and its frequency and amplitude (Separation, n = 5 cells from 3 mice, Cx43 OE, n = 4 cells from 3 mice). (j) Heatmap and averaged F/F traces of LHb neuronal Ca^2+^ responses to stimulation, with quantification of the area under the curve (AUC) during *in vivo* fiber photometry recordings (Separation n = 9 stimulation form 3 mice, Cx43 OE, n = 18 stimulation from 6 mice).

We next examined whether Cx43 restoration normalised astrocytic Ca^2+^ dynamics. *In vivo* photometry recording of LHb astrocytes showed that overexpression of astrocytic Cx43 markedly reduced Ca^2+^signals in mice with parental separation (Fig. 7b), indicating recovery of astrocytic Ca^2+^ homeostasis.

Because Cx43 restoration normalised astrocytic Ca^2+^ activity, we examined whether it also restored the Ca^2+^-Rac1–p-Ezrin cytoskeletal pathway implicated in astrocytic leaflet remodelling. Parental separation-induced reduction in phosphorylated Ezrin (p-Ezrin) was significantly restored following astrocytic Cx43 overexpression (Fig. 7c). Subsequently, we assessed whether the recovery of this pathway was accompanied by restoration of astrocytic morphology. The Cx43 overexpression significantly increased the volume fraction of leaflets by 7.36% ± 0.84% (P < 0.0001) in the Separation group (Fig. 7d). Sholl analysis of LHb astrocytes showed that their complexity was also significantly increased in the Separation group following Cx43 overexpression (Extended Data Fig. 8d), accompanied by a restored ALDH1L1-RFP+ astrocyte cell domains (increased by 55.76% ± 17.13%, n = 5, P < 0.05, Extended Data Fig. 8d).

We next investigated whether recovery of astrocytic structural complexity recovered astrocyte–synapse interactions. Super-resolution imaging of astrocytes and synaptic markers showed that overexpression of Cx43 in the LHb astrocytes restored the co-localisation of ALDH1L1-RFP with the excitatory synaptic markers VGLUT2 and Homer1 increased by 33.64% ± 7.79% in the Separation group (Separation n = 5, Cx43 overexpression, n = 4, P < 0.01) (Fig. 7e, Extended Data Fig. 8e). The decreased density of VGLUT2 or Homer1 puncta connected with individual astrocytes in the Separation group was also restored following Cx43 overexpression (VGLUT2: increased by 48.47% ± 13.97%, P < 0.05, Homer1: increased by 28.64% ± 11.44%, Separation n = 5, Cx43 overexpression, n = 4, P < 0.05), without affecting the total VGLUT2 and Homer1 co-localisation suggestive of the intact excitatory synapses (Extended Data Fig. 8e). The deficits in inhibitory synapses in the LHb were rescued as demonstrated by significantly increased co-localisation of ALDH1L1-RFP with the inhibitory synaptic markers Synapsin and Gephyrin in the Separation group following Cx43 overexpression (increased by 150.8% ± 33.80%, Separation n = 5, Cx43 overexpression, n = 5, P < 0.01) (Fig. 7f, Extended Data Fig. 7f). The reduction in inhibitory synapses (Synapsin or Gephyrin) puncta associated with individual astrocytes in the Separation group was also reversed (Extended Data Fig. 8f, Synapsin: increased by 99.58% ± 19.19%, Separation n = 5, Cx43 overexpression, n = 4, P < 0.001, Gephyrin: increased by 136.5% ± 24.90%, Separation n = 5, Cx43 overexpression, n = 4, P < 0.001). Of note, while Cx43 overexpression rescued astrocyte-associated inhibitory synapses, it did not alter the overall total Synapsin and Gephyrin co-localisation in the LHb (Extended Data Fig. 8f). Furthermore, TEM revealed corresponding ultrastructural changes where astrocytic Cx43 overexpression normalised parental separation-induced increase in the gap area between astrocytes and synapses (Fig. 7g, decreased by 866.2 ± 307.5 nm^2^, Separation n = 17 cells from 4 mice; Cx43 overexpression, n = 14 cells from 4 mice, P < 0.01), demonstrating a rescue of the multipartite synapse ultrastructure. The perimeter of neuronal boutons and spines was unchanged, indicating an unaffected pre- and post-synaptic neuronal compartment (Fig. 7g).

We next examined whether structural recovery of astrocyte–synapse interactions translated into functional normalisation of LHb. Neuronal whole-cell patch clamp recordings revealed that Cx43 overexpression in astrocytes in the Separation group significantly increased the frequency, but not the amplitude, of mEPSC (Fig. 7h) and mIPSC (Fig. 7i), suggesting normalised synaptic transmission in mice subjected to early-life parental separation. Consistent with improved synaptic function, neuronal Ca^2+^ photometry revealed that the excessive LHb neuronal activity induced by parental separation was mitigated following astrocytic Cx43 restoration (Fig. 7j).

Finally, we explored whether the functional normalisation of LHb induced by Cx43 overexpression can improve behavioural abnormalities. As expected, mice in the Separation group with Cx43 overexpression in the LHb astrocytes exhibited significantly reduced swimming time in the FST compared to mice in the Separation group without Cx43 overexpression (Fig. 7k), indicating alleviated depression-like behaviour. However, the distance travelled in the centre zone of the OFT remained unaffected (Extended Data Fig. 8g), suggesting that anxiety-like behaviour was not linked to Cx43 level in the LHb.

Collectively, these findings demonstrate that restoring astrocytic Cx43 reverses the stress-induced pathological cascade, normalising astrocytic Ca^2+^ signalling and p-Ezrin-dependent cytoskeletal organisation, restoring astrocytic morphology and tripartite synapse architecture, reducing LHb hyperactivity and alleviating depressive-like behaviours. These results identify astrocytic Cx43-dependent network homeostasis as a reversible mechanism underlying adolescent stress-induced depression.

## Discussion

Our study shifts the framework from a neuron-dominant model of LHb dysfunction to a glial-neuronal network model, in which astrocytic Cx43 coupling maintains synaptic architecture while Cx43 downregulation initiates the pathological cascade leading to adolescent depression. We identified a previously unrecognised astrocytic mechanism through which early-life stress disrupts LHb circuit and drives adolescent depressive-like behaviour. LHb astrocytes demonstrate regional specialisation, which is selectively compromised by parental separation by a Cx43-dependent mechanism acting through Rac1–p-Ezrin cytoskeletal pathway. This remodelling disrupts astrocyte engagement with both excitatory and inhibitory synapses, and leads to LHb neuronal hyperactivity. Both suppression of excessive astrocytic Ca^2+^ signalling and restoration of astrocytic Cx43 reverse this pathological cascade. Together, these establish a glial-neuronal network model, in which astrocytic Cx43-dependent homeostasis preserves tripartite synapse architecture and represents a reversible mechanism underlying adolescent stress-induced depression.

Astrocytes are increasingly recognised as regionally specialised regulators of neural circuit function ^9,55,56^. Consistent with this view, our study identifies the LHb as a region containing distinct astrocytes, characterised by compact domains, less complex primary branching, relatively enriched leaflets, and high Cx43 levels. This regional specialisation creates a Cx43-dependent astrocytic network vulnerable to early-life stress. Astrocytic malfunction has been implicated in the pathogenesis of depression and stress-induced depressive behaviours ^13,14^. Astrocyte ablation from the mPFC ^13,57^, disruption of astrocytic homeostatic pathways including glutamate transporters or connexins ^32,58^, and impaired glutamate clearance ^59–62^ can produce or be associated with depressive-like phenotypes ^63^. We identify the LHb as a previously underappreciated site of region-specific astrocytic vulnerability, where disruption of Cx43-dependent astrocytic network homeostasis is sufficient to drive local circuit dysfunction and recapitulate depressive-like behaviour of mice with early life parental separation. Specifically, parental separation reduced astrocytic Cx43 and gap-junction coupling, leading to excessive astrocytic Ca^2+^ activity, Rac1-GTPase-dependent cytoskeletal remodelling regulated by p-Ezrin ^64–67^, astrocytic atrophy and leaflet retraction. Astrocyte atrophy impaired their engagement with neuronal synapses, consistent with the role of perisynaptic astrocytic leaflets in multipartite synapse organisation ^40,68,69^, resulting in reduced excitatory and inhibitory synaptic transmission and aberrant LHb neuronal activity. Incidentally, connexins have recently emerged as potential therapeutic targets in depression ^70–72^. Our data show that astrocytic Cx43 loss in the LHb is sufficient to phenocopy the stress-induced structural, synaptic, neuronal, and behavioural abnormalities, whereas astrocytic Cx43 restoration rescues these pathological changes. Together with evidence that astrocytic atrophy occurs in rodent models and human depression ^67,73,74^, these findings identify Cx43-dependent LHb astrocytopathy as a reversible mechanism contributing to stress-induced adolescent depression.

A key conceptual advance of our study lies in establishing direct link between astrocytic atrophy and LHb synaptic and circuit malfunction. Astrocytic leaflets form a critical structural component of tripartite synapse architecture. Loss of leaflets after parental separation or astrocytic Cx43 suppression reduced astrocyte engagement with excitatory and inhibitory synapses without reducing total synaptic numbers, indicating that stress disrupts the functional integration of astrocytes with synaptic units rather than globally eliminating neuronal synapses. This astrocyte–synapse disconnection was accompanied by reduced miniature excitatory and inhibitory postsynaptic currents and, unexpectedly, with elevated LHb neuronal Ca^2+^. This differs from adult depression models, in which LHb hyperactivity has often been attributed to enhanced excitatory transmission, increased glutamatergic drive or altered excitatory/inhibitory balance^75–77^. The discrepancy may reflect developmental stage-dependent mechanisms, as adolescence is marked by ongoing synaptic maturation, inhibitory circuit refinement and dynamic regulation of intrinsic excitability^78,79^. Thus, in adolescent stress, disruption of Cx43-dependent astrocytic support may weaken synaptic integration while contributing to aberrant neuronal hyperactivity. Conversely, Cx43 restoration rebuilt leaflet complexity, re-established astrocyte-associated excitatory and inhibitory synaptic structures, normalised astrocyte–synapse ultrastructural relationships, improved synaptic transmission and reduced LHb hyperactivity. These findings indicate that astrocytic morphology is not merely a marker of cellular stress but an active structural determinant of synaptic integrity and circuit output in the LHb.

Our study identified a molecular pathway linking deficiency in Cx43 to astrocytic structural remodelling. Loss of astrocytic Cx43 increased spontaneous Ca^2+^ activity in LHb astrocytes, contrasting with our previous findings in hippocampal astrocytes, where early-life stress suppressed astrocytic Ca^2+^ signalling ^33^. This regional divergence highlights the idea that astrocyte responses to stress are brain-region specific and may be central to LHb vulnerability. Mechanistically, increased astrocytic Ca^2+^ activity activates Rac1, a regulator of astrocytic process growth and cytoskeletal organisation ^49,80,81^, which in turn reduces p-Ezrin, a membrane–actin linker involved in astrocyte–synapse interactions and fine-process architecture ^82,83,84^. Because variations in astrocytic Ezrin expression and phosphorylation have been linked to stress resilience through regulation of astrocyte structure^85,66,86^, our findings place Ezrin downstream of a Cx43-Ca^2+^-Rac1 cascade in the LHb. This pathway provides a mechanistic explanation for how impaired astrocytic network communication is converted into leaflet retraction, tripartite synapse disruption and circuit dysfunction during adolescent stress.

This study has several limitations. We identify LHb astrocytes as a regionally specialised and stress-sensitive, but whether similar or divergent Cx43-dependent mechanisms operate across other depression-relevant brain regions remains unresolved. Moreover, although our data establish causality in a mouse model of early-life stress, whether comparable astrocytic Cx43-dependent pathology occurs in human adolescent depression requires further investigation. Nonetheless, our study identifies astrocytic Cx43-dependent network homeostasis as a key determinant of LHb circuit resilience during adolescence and reveals a reversible glial mechanism by which regionally specialised astrocytes regulate vulnerability to stress-induced depressive pathology.

## Conclusion

We identified astrocytic Cx43-dependent network failure as a previously unrecognised and reversible mechanism linking early-life stress to LHb circuit malfunction and adolescent depressive-like behaviour. Stress-induced loss of astrocytic Cx43 disrupts gap-junction coupling, drives aberrant astrocytic Ca²□hyperactivity, activates the Rac1-p-Ezrin cytoskeletal pathway and triggers astrocytic leaflet retraction and cellular atrophy. Glial remodelling compromises tripartite synapse integrity, weakens synaptic transmission and uncouples synaptic input from LHb neuronal output, resulting in maladaptive LHb hyperactivity. Our findings establish LHb astrocytic Cx43 as a causal regulator of stress vulnerability and a potential therapeutic entry point for adolescent depression.

## Materials and methods

### Animal model establishment

Animal experiments were approved by the Animal Care and Use Committee (Approval# SYSU-IACUC-2022-002016) of Sun Yat-Sen University Institutional and performed following the National Institute of Health Guide for the Care and Use of Laboratory Animals. C57BL/6J mice were housed on a 12 h/12 h light/dark cycle at 22 ± 1□and a humidity of 50% ± 10%. The model of parental separation was established as previously described ^87^. Briefly, pups of both sexes were randomly divided into two groups. Parental separation occurred from postnatal day (P)2-P12 (equivalent to human infancy ^81^, when pups were separated from their parents and housed individually from 9:00-17:00 daily (Separation group) with soft bedding in small boxes. Another group of mice underwent a sire-only condition to control for milk availability, which was separated from their dams but stayed with their sires during the daily 9:00-17:00 separation period from P2-P12 (Control group). Mice in the Separation group were sacrificed at P23, while mice in the Control group (Fig. 1) were sacrificed at P10, P23, or 2 months old.

### AAV injection

To conditionally overexpress Cx43 in the LHb, the mouse *Gja1* coding sequence (NM_010288.4) along with a P2A sequence was cloned into the pAAV-hALDH1L1-P2A-RFP, which was then packaged into the AAV2/5 virus (AAV2/5-ALDH1L1-Gja1-P2A-RFP, Weizhen Biology, Jinan, China). For knockdown of Cx43, shRNA targeting Gja1 was inserted into pAAV-U6-shRNA-CMV-Intron-RFP and packaged into AAV2/5 (AAV2/5-ALDH1L1-shRNA-Gja1-RFP, Weizhen Biology, Jinan, China). For stereotaxic surgery, P6 mice were anaesthetised with isoflurane (2-3%) and AAV (200 nL, titer: 1.5-6.5 x 10^12^ vg/μL) was injected into the LHb -0.7 x 1.72 mm anteroposterior from the bregma, ± 0.4 mm medial-lateral, -2.5 mm dorsoventral. To specifically express GCaMP in astrocytes and neurons, AAV2/5-gfaABC1D-GCaMP6s-WPRE-pA (100 nL, titer: 1 x 10^13^ vg/μL, Taitool, S0530) and AAV2/5-hSyn-GCaMP6s-WPRE-pA (100 nL, titer: 1 x 10^13^ vg/Μl, Taitool, S0225) were injected into mice LHb. CalEx was packaged into AAV (AAV5-pZac2.1-GfaABC1D-mCherry-hPMCA2w/b, titer: 8.04x10^13^/muL) and injected into LHb to inhibit astrocytic intracellular Ca^2+^ *in vivo*.

### Behavioural Tests

Three days before the test (P20), mice were allowed to familiarise themselves with the test arena and handled by the experimenter daily. All mice were tested from 9:00 to 17:00, and the recordings were analysed using VisuTrack software (Xinruan, Shanghai, China). For the Open Field Test, mice were placed in the centre of the open field arena (50 x 50 x 50 cm) and recorded for 20 min. The total distance and percentage of distance in the central area were analysed. For the Forced Swim Test, the same mice were placed in the water (25 □) for 7 min and recorded. VisuTrack software was used to distinguish ‘swimming’ or ‘struggling’ status according to their activity.

### *In vivo* fibre photometry

Mice received AAV injections to genetically expressed Ca^2+^ indicator GCaMP specifically in LHb at P6. Three weeks after AAV expression, the fibre (Ø1.25mm ferrule, 0.37 NA, 3.0 mm length) were implanted into 300 μm above LHb. 5 days after implantation, the signal of GCaMP was recorded using Multi-Channel Fiber Photometry Device (Inper Ltd., China). The light power was approximately 35 μW for the 470 nm wavelength and the 15 μW for the 410 nm wavelength at the tip of optic fibre. During recording, mice were placed in a 25 x 25 cm chamber that permitted free movement, and clamping was randomly occurred during recording. Data were processed and analysed using Inper Data Analysis software.

### Acute brain slices

Acute brain slices were prepared following the previously published protocols ^88^. Fresh brains from P23-P25 pups were incubated in cool NMDG-HEPES artificial cerebrospinal fluid (aCSF) for 1 min before slicing. Brain slices were then transferred into 34□NMDG-HEPES aCSF, and Na^+^ spike-in solution was supplied every 2 min for 10 min. Afterwards, the brain slices were incubated in HEPES aCSF for 1 h before experiments in oxygenated (95% O_2_, 5% CO_2_) recording aCSF.

### Calcium imaging

Acute brain slices were incubated with 5 μM Fluo4-AM solution for 40 min at 34□, washed out for 5 min, then incubated with 100 μM 2-APB (A106408) for 30 min or directly transferred into a recording chamber under an FV3000 confocal microscope (Olympus) when perfused with oxygenated recording aCSF. For Ca^2+^ imaging, the fluorescence signal of Fluo4-AM was collected from ALDH1L1-RFP positive cells every 2.17 s for 150 frames in total and quantified using FIJI (NIH). The average fluorescence intensity of the first 10 frames served as the baseline (F_0_), against which the fluorescence intensity of each frame (F_t_) was normalised (dF/F_0_). A de-trend and baseline correction pre-treatment of the dF/F_0_ was performed using PyCharm to remove the trend caused by fluorescence decline of the whole slice or z-plane drift and noise. Then, the events per minute, amplitude, and area under curve (AUC) were calculated.

### Immunofluorescence staining

The brains were dehydrated in 30% sucrose and embedded in OCT for sectioning (16 μm). Sections were incubated in 1% Triton-X100 in PBS (Triton-PBS), blocked with 2% bovine serum albumin (BSA), and then incubated with primary antibodies followed by secondary antibodies at room temperature for 2 h and coverslipped. The primary antibodies included: Rabbit anti-BLBP (BLBP, ab32423, Abcam), Goat anti-Glial Fibrillary Acidic Protein (GFAP, ab53554, Abcam), Rabbit anti-Aldehyde Dehydrogenase 1 Family Member L1 (ALDH1L1, ab87117, Abcam), Rat anti-GFP (OB-PRT018-02, Oasis biofarm), Rat anti-RFP (OB-PRT017-01, Oasis biofarm), Rabbit anti-Cx43 (C6219, Sigma), Rabbit anti-Cx30 (ab200866, Abcam), Goat anti-Ionised calcium-binding adapter molecule 1 (Iba1, ab5076, Abcam), Rabbit anti-APC (CC1, OB-PRB070-01, Oasis biofarm), Goat anti-Platelet-Derived Growth Factor Receptor Alpha (PDGFRα, AF1062, R&D System), Guinea pig anti-Synapsin (106004, Synaptic System), Guinea pig anti-Vesicular Glutamate Transporter 2 (VGLTU2, AB2251-1, Millipore), Rabbit anti-Gephyrin (12681, Proteintech), Rabbit anti-Homer Scaffold Protein 1 (Homer1, 160003, Synaptic System) Rabbit anti p-Ezirn (PA5-37763, Invitrogen), Mouse anti active Rac1 (26903, Neweast biosciences) Secondary antibodies included: Donkey anti-Rabbit Alexa Fluor 568 (A10042, Invitrogen), Donkey anti-Rabbit Alexa Fluor 647 (A32795, Invitrogen), Donkey anti-Goat Alexa Fluor 488 (A32814, Invitrogen), Donkey anti-Goat Alexa Fluor 568 (A11057, Invitrogen), Goat anti-Guinea pig Alexa Fluor 647 (A21450, Invitrogen), Donkey anti-Rat Alexa Fluor 488 (A48269, Invitrogen), Donkey anti-Rat Alexa Fluor 594 (A21209, Invitrogen). Sections were imaged by a VS200 Research Slide Scanner (Olympus), an FV3000 confocal microscope (Olympus), and Ixplore SpinSR confocal microscope (Olympus). Super-resolution images of connexins were captured using an Ixplore SpinSR confocal super-resolution system with a UPLAPO60XOHR objective (Olympus). Super-resolution images of the multipartite synapse were captured by the LIGHTNING module of Leica Stellaris5 confocal. For multipartite synapic structure analysis, 3D reconstruction of synaptic marker puncta and ALDH1L1-RFP positive astrocytes was carried out in Imaris 9.9 first. Subsequently, synaptic puncta located within a distance less than their own diameter from ALDH1L1-RFP were identified as ALDH1L1-RFP connected puncta, and such connection was referred to as ‘co-localisation’.

### Gap Junction Fluorescence Recovery After Photobleaching (gap-FRAP) assay

Acute brain slices were incubated in 10 µM Calcein for 30 min at 34□before being transferred into the recording chamber under an FV3000 confocal microscope (Olympus). Baseline fluorescence was recorded at a low-power 488 nm laser excitation every 5s for 10 cycles over a 158 x 158 µm field. This was followed by photobleaching of the target astrocyte using higher laser power over a 5 x 5 µm field for 20 s. Recording continued for 10 min post-bleaching to capture fluorescence recovery. The fluorescence intensity at each time point was normalised to F0 (the average baseline fluorescence intensity) and expressed as a percentage of F0 (Ft/F0).

### Dye uptake

Dye uptake in acute brain slices was measured following a previously published protocol ^29^. Briefly, acute brain slices were incubated in aCSF with or without hemichannel blocker CBX (200 µM), for 15 min at room temperature, followed by EtBr (4 µM) for 10 min at room temperature. Then, brain slices were fixed with 4% paraformaldehyde for 1 h before immunostaining. For immunofluorescence staining, slices were incubated with 1% Triton-PBS for 30 min, followed by 2% BSA blocking. Then slices were incubated with primary antibodies (Rat anti-GFP, OB-PRT018-02, Oasis biofarm) at 4□overnight, followed by a secondary antibody (Donkey anti-Rat Alexa Fluor 647, A48269, Invitrogen). Consecutive images were collected from the same depth below the surface (about 10 µm) of the slices with the same interval for LHb using an Ixplore SpinSR confocal microscope (Olympus).

### Single-cell RNA-sequencing

LHb tissue was collected at P23 for single-cell RNA sequencing. For each condition, LHb tissue from more than ten mice was pooled before library preparation, generating one pooled single-cell dataset for the control condition and one for the maternal separation condition.

Minced tissues (∼2–4 mm³) were enzymatically dissociated in the enzymatic solution. The resulting single-cell suspensions were filtered through 70-μm and 30-μm cell strainers (Miltenyi Biotech, PN: 130-098-458 and 130-098-462), followed by red blood cell removal using Red Blood Cell Lysis Solution (Miltenyi Biotech, PN: 130-094-183). Cell viability was assessed using Countstar Rigel (Alit Biotech), and dead cells were removed using the Dead Cell Removal Kit (Miltenyi Biotech, PN: 130-090-101) when required. Cells were resuspended in 1× PBS containing 0.04% BSA at a final concentration of 700–1,200 cells/μL and loaded onto the 10x Chromium platform. Single-cell libraries were generated using the 10x Chromium Single Cell 3′ Kit according to the manufacturer’s instructions and sequenced on an Illumina NovaSeq platform with 150-bp paired-end reads.

Single-cell RNA-sequencing data were analysed in R v.4.5.0 using Seurat v.5.3.1. Gene–cell count matrices from control and separation samples were imported separately and combined after quality control. Cells were filtered on the basis of the number of detected genes, total transcript counts and mitochondrial transcript fraction. The filtered datasets were merged into a single Seurat object for downstream analysis. Data were normalised using SCTransform v2. Highly variable features were used for principal-component analysis (PCA), and 50 principal components were calculated. The first 30 principal components were used for nearest-neighbor graph construction, dimensionality reduction and global clustering. Graph-based clustering was performed at a resolution of 0.4, yielding 24 transcriptionally defined clusters. Cell identities were assigned on the basis of established lineage markers together with cluster-specific gene-expression patterns. Major populations included astrocytes, oligodendrocytes, oligodendrocyte precursor cells, microglia, macrophages, other immune cells, neurons, endothelial cells, pericytes, smooth muscle cells, fibroblasts, ependymal cells, choroid plexus epithelial cells and pinealocytes. Representative marker expression was visualised using feature plots and dot plots. Cluster-specific markers were identified using the Wilcoxon rank-sum test implemented in Seurat. Cells annotated as astrocytes were extracted for astrocyte-specific analysis.

Cell–cell communication was inferred using CellChat v.2.2.0.9001. Control and separation datasets were analysed independently. Overall astrocyte-to-neuron communication was quantified from the aggregated CellChat interaction-weight matrix. Pathway-specific networks were obtained from pathway-level communication probabilities. For comparison of control and separation networks, identical node sizes were used across conditions to prevent differences in cell abundance from influencing network visualisation. For each signalling pathway, control and separation circle plots were plotted using the same maximum edge weight and edge-width scale.

### Slice Culture

Before slicing, the slice medium (50% DMEM, 25% HBSS, and 1% penicillin-streptomycin) and culture medium (50% DMEM, 25% HBSS, 25% heat-inactivated horse serum, 1% N2 and penicillin-streptomycin) were prepared. Slice culture followed a previously published protocol ^89^. Briefly, mice at P23 were sacrificed, and the brain was assessed. The meninges were removed, and the brain was coronally cut to 300 μm in slice medium. The slice containing LHb was collected and transferred to 0.4 μm Millicell-CM organotypic culture inserts (Millipore, Cat No. PICM0RG50). Slice were then cultured at 37 °C and 5% CO_2_ for 12 h in culture medium, then fixed with 4% PFA and mounted for imaging.

### Electron microscopy

Mice were perfused with 0.1 M phosphate buffer, followed by a fixative containing 4% paraformaldehyde, 0.2 M phosphate buffer, and 25% glutaraldehyde 4□for at least 48 h. The tissues were then incubated with OsO4 for 2 h and immersed in a graded acetone series to dehydrate, followed by embedding in Epon. The Semithin and ultrathin sections were prepared and viewed under a transmission electron microscope (TECNAI10, Philips). Astrocytes were identified by their relatively clear cytoplasm and presence of glycogen granules and bundles of filaments. Synapses were identified by the boutons filled with small vesicles and adjacent dendritic spines, where the postsynapse appeared with darkened band, indicating the postsynaptic density area ^45–48^. The astrocyte processes wrapping synapses were identified as peri-synaptic astrocyte processes (PAPs), and the area between PAPs and synaptic cleft was measured and represented as “gap area”.

### Patch-based dye injection into astrocytes

The patch-based dye infusion was performed following a previously published method ^90^. Briefly, slices were perfused with oxygenated aCSF. Sulforhodamine B (SRB) was diluted to a concentration of 1 mg/mL in the pipette internal solution, which contained 140 mM D-gluconic acid, 10 mM KCl, 1 mM MgCl_2_, 10 mM HEPES, 11 mM EGTA, 1.5 mM Mg-ATP, 0.3 mM Li-ATP, with pH adjusted to 7.3 with KOH. Cells were loaded with SRB and allowed to diffuse for 40 min under current-clamp mode. The slices were then imaged using an Ixplore SpinSR confocal microscope (Olympus) immediately after the diffusion time ended.

### Whole-cell patch-clamp

Brain slices were transferred into the recording chamber and perfused with oxygenated aCSF. Then, neurons surrounding RFP-positive astrocytes were identified and selected for recording. For miniature excitatory postsynaptic current (mEPSC) recording, cells were voltage clamped at -60 mV using an internal solution containing (in mM): 123 K-gluconate, 12 KCl, 10 HEPES, 0.2 EGTA, 4 Mg-ATP, 0.3 Na-GTP, 10 Na_2_-phosphocreatine. Tetrodotoxin and bicuculline were applied to isolate excitatory currents. For miniature inhibitory postsynaptic current (mIPSC) recording, cells were voltage clamped at -70 mV with an internal solution containing (in mM): 140 KCl, 4 MgCl_2_, 0.1 EGTA, 10 HEPES, 0.3 Na_2_GTP, 2 Na_2_ATP. Tetrodotoxin and NBQX were applied to isolate inhibitory currents. Using Clampfit 10.7 to calculate the frequency and amplitude of each cell, and each recorded currents was extracted to make an average current, further reflecting the amplitude of different groups.

### Primary astrocyte culture and AAV infection

Astrocytes culture medium (10% FBS, 1% penicillin-streptomycin solution, 1% GlutaMax in DMEM), papain solution (1X EBSS 9.78 mL, 100 mM MgSO_4_ 100 μL, 45% Glucose 80 μL, 0.5 M EGTA 40 μL), papain buffer (L-cysteine 5 mg, papain 10 mg, DNase 300 μL in 10 mL papain solution), 6 x Hi-ovo (Ovomucoid 1g, BSA 1 g, DPBS (with Ca^2+^/Mg^2+^) 16.7 mL), Low-ovo (250 μL 6x Hi-ovo, 9.45 mL DPBS (with Ca^2+^/Mg^2+^), 300 μL DNase) were prepared.

Mice were sacrificed at P6, and the LHb was isolated by microdissection in pre-cooled DPBS (without Ca^2+^/Mg^2+^). Then, the tissue was digested in papain buffer at 37□ for 1 h, and then mechanically dissociated by trituration with a pipette. The supernatant was removed before adding the Low-ovo to resuspend the tissue. The supernatant was transferred into 1x Hi-ovo and centrifuged at 1500 RPM for 5 min. The resulting cell pellet was resuspended with astrocyte culture medium and then seeded onto a 6 cm dish.

3 h before AAV infection, the culture medium was replaced with infection culture medium (2% FBS, 1%, penicillin-streptomycin solution, 2% B27). AAV (1.5 × 10^10^ vg/μL) was added for 16 h before replacing the medium with astrocyte culture medium for 3 days before protein extraction.

### Protein extraction and western blotting

RIPA lysis buffer (50 μL, with phosphatase inhibitor and protease and phosphatase inhibitor) was added to each well for cell lysis. The solution was mixed with an equal volume of SDS-PAGE and heated at 100□for 5 min. Then, 10 μg samples were used for electrophoresis, and the separated proteins were transferred to a polyvinylidene difluoride (PVDF) membrane. The membrane was incubated with primary antibodies at 4□, followed by a secondary antibody at room temperature for 2 h. The Amersham ECL Prime Western Blotting Detection Reagent (Cytiva) and ChampChemi 610 Chemiluminescence Imaging System (Beijing Sage Creation, China) were used to detect the bands.

### Statistical analysis

Data are presented as mean ±□standard error of the mean (SEM) and were analysed using GraphPad Prism (V9.0). An unpaired t-test was used to compare two groups, whereas one-way ANOVA followed by Tukey’s post hoc test was used for comparisons involving more than two groups. Sholl-analysis data were analysed using two-way ANOVA followed by Tukey’s test. P < 0.05 was considered statistically significant.

## Acknowledgement

This work was supported by grants from the National Natural Science Foundation of China (32671294 to C.Y.; W2511095 to J.N. and A.V.; 32271034 to J.N.; 82260272 to D.L.; 82201707 to X.H.), Shenzhen Fundamental Research Program (RCJC20231211090018040 and ZDSYS20220606100801003 to C.Y.; JCYJ20210324121214039 and RCBS20210706092411028 to Y.S.; JCYJ20220530144816038 and RCBS20221008093118042 to Q.W.; JCYJ20240813150421029 to X.H.), Chongqing Natural Science Fund for Distinguished Young Scholars (CSTB 2023NSCQ-JQX0030 to J.N.), National Key Research and Development Program of China (2021ZD0201703 to J.N.), Guangdong Basic and Applied Basic Research Foundation (2023A1515010817 to Y.S.), Science and Technology Planning Project of Guangdong Province (2021A1515110268 to Y.S.), Shenzhen Medical Research Fund (A2303014 to Y.S.), and “3510” Medical Innovation Leadership Program, Fundamental Research Funds for the Central Universities, Sun Yat-sen University (20263510JCQN011 to C.Y.). AV was supported by 2023 Key Support Project of the Liaoning Provincial Department of Science and Technology to Support the High-Quality Development of China Medical University ([2023]61-7), and by Slovenian Research Agency grant J4-60077.

## Conflict of interest statement

The authors declare no conflict of interest.

## Data availability

All data that support the findings of this study are available from the corresponding author upon reasonable request.

## Extended Figures

**Extended Fig 1.**
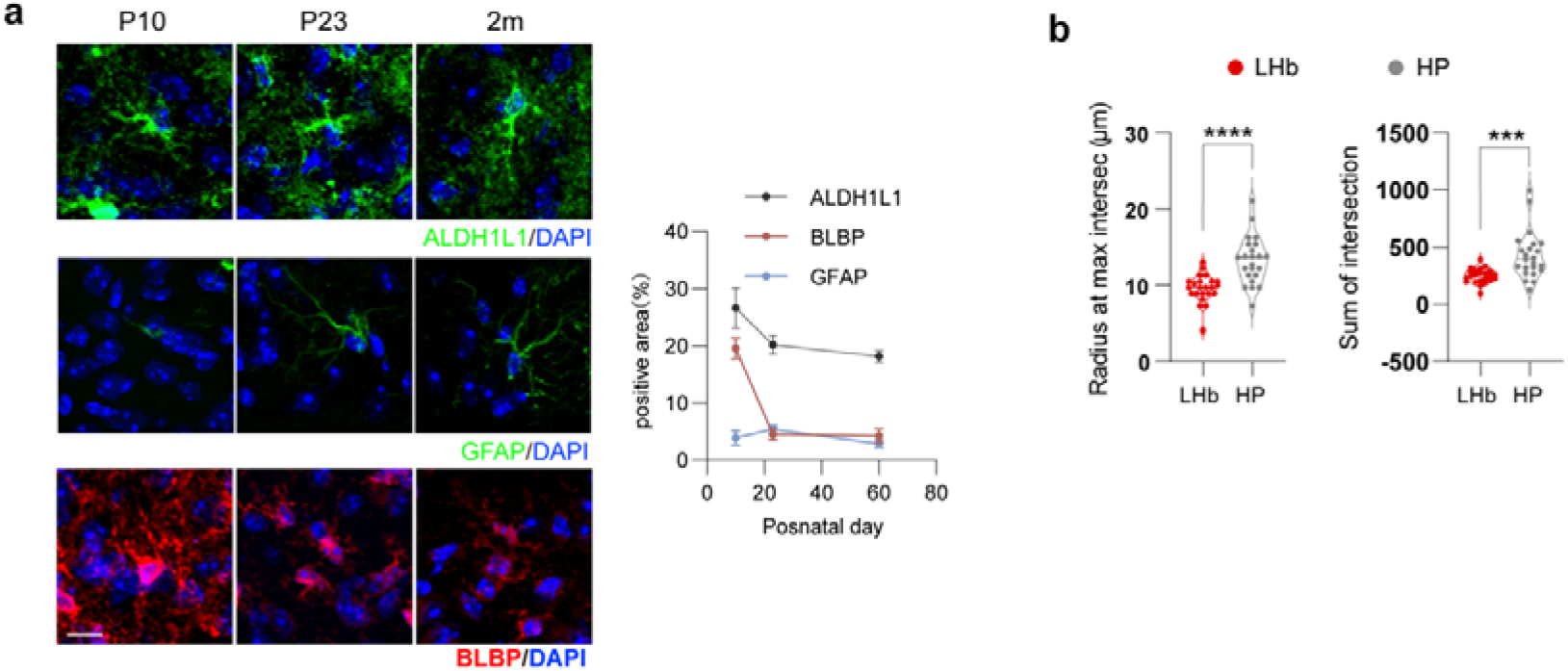
Features of astrocytic morphology and its marker expression in LHb. (a) Representative images of BLBP and GFAP staining and quantification in the LHb at postnatal day (P)10, P23, and 2-month-old (P60) and their quantification (Scale bar, 10 μm). (b) Quantification of the radius at max intersection and the sum of intersection in Sholl analysis.

**Extended Fig 2.**
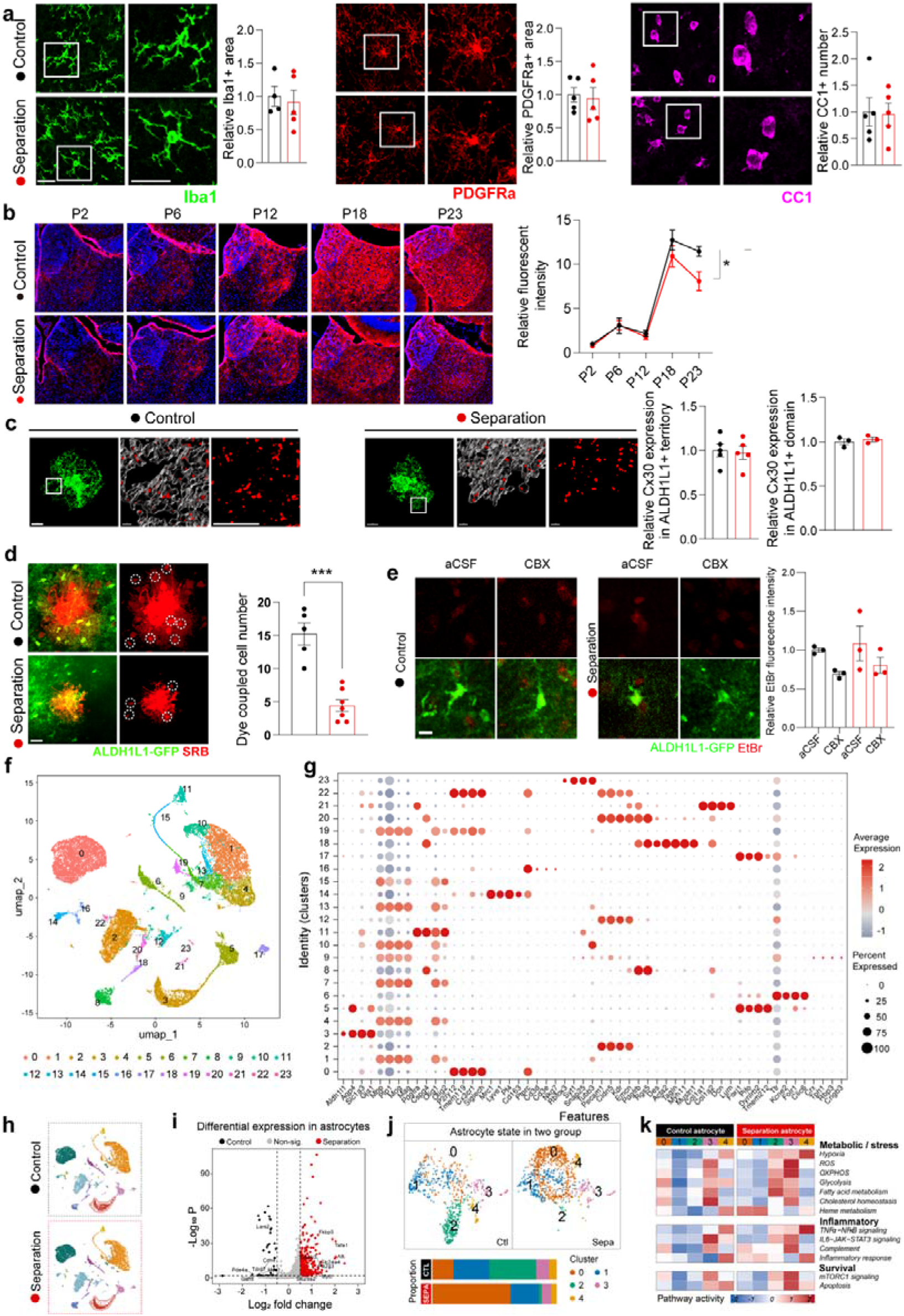
Neuroglial cells in the LHb, Cx expression and its function in the parental separation model. (a) Representative images of Iba1, PDGFRa, and CC1 staining in the LHb, and quantification of the areas (Iba1, Control n = 4 mice, Separation n = 5, PDGFRa and CC1 n = 5 mice; Scale bar, 20 μm). (b) Representative images of Cx43 at different stage during development and the quantification of Cx43 fluorescence intensity (n = 3 mice, Scale bar, 100 μm). (c) Representative images of Cx30 immunostaining and ALDH1L1-GFP fluorescence in the LHb, and the quantification of Cx30 in ALDH1L1-positive territory and domain (territory n = 5 mice, domain n = 3 mice; Scale bar, 5 μm). (d) Fluorescent images of SRB in ALDH1L1-GFP labelled cells in LHb, and quantification of dye coupled cells number (Control n = 5 cells from 3 mice, Separation n = 7 cells from 3 mice; Scale bar, 20 μm). (e) Representative image of EtBr and ALDH1L1-GFP staining and quantification of EtBr uptake by ALDH1L1-GFP labelled astrocytes (n = 3 mice, Scale bar, 10 μm). (f) UMAP representation of the LHB single-cell RNA-sequencing dataset showing 24 transcriptionally defined clusters (clusters 0–23) before cell-type annotation. (g) Dot plot showing the expression of representative canonical marker genes across the 24 clusters used to support cell-type annotation. Dot size indicates the percentage of cells expressing each gene, and colour indicates scaled average expression. (h) Feature plots showing representative marker genes used to support cell-type annotation. (i) UMAP representation of LHb cells separated by control and maternal separation condition. (j) Differential gene-expression analysis of astrocytes from control and separation conditions. (k) Astrocyte-specific UMAPs showing five transcriptional states in control and separation conditions. The stacked bar plot shows the relative representation of the five subtypes within each condition.

**Extended Fig 3.**
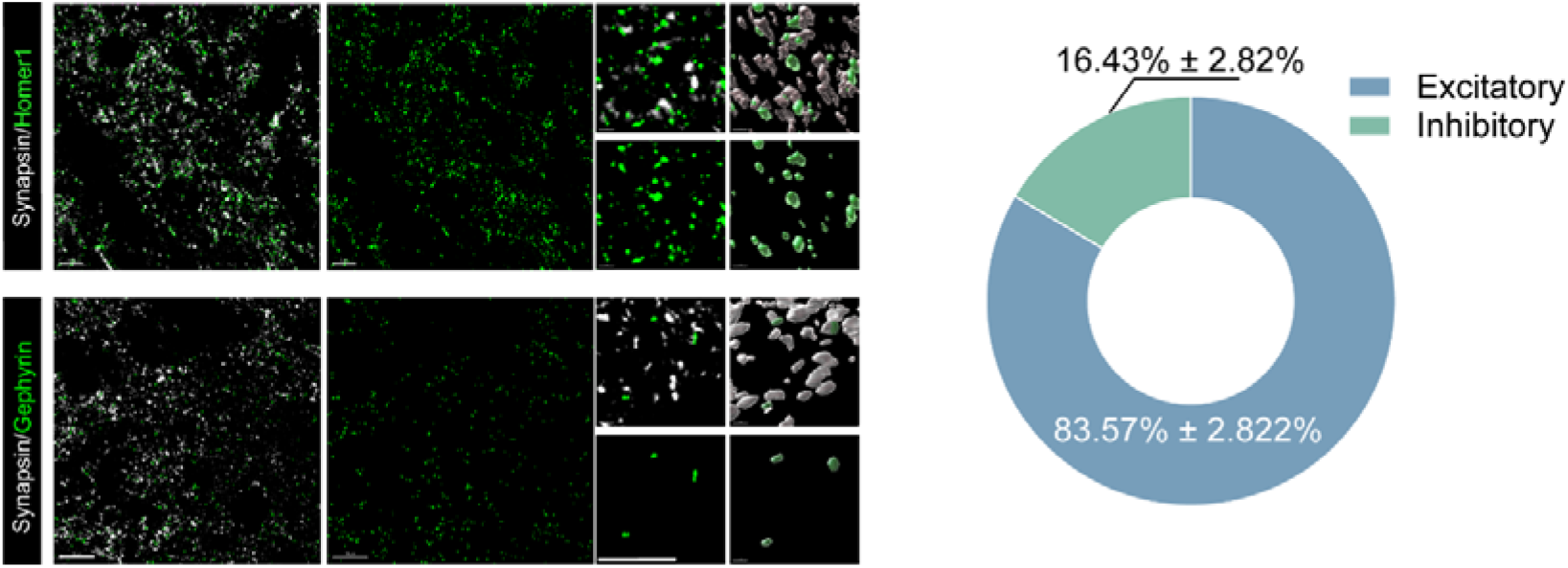
The percentage of excitatory and inhibitory synapses in LHb. Representative images of Synapsin, Homer1, and Gephyrin in the LHb, and the percentage of Synapsin co-localised with Homer1 (Excitatory) or Gephyrin (Inhibitory), n = 3 mice; Scale bar, left, 10 μm, right, 5 μm.

**Extended Fig 4.**
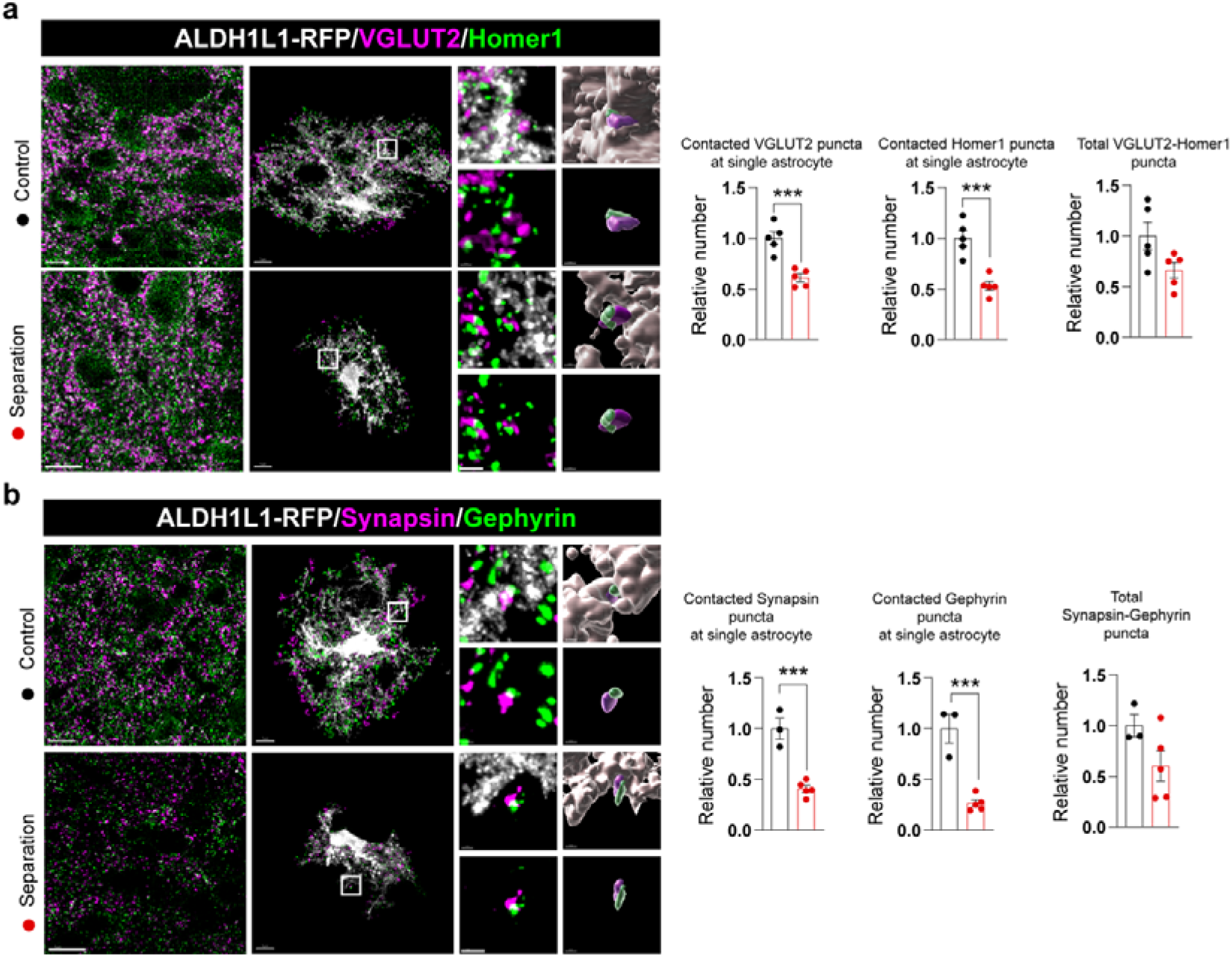
Astrocytes mediated remodelling of tripartite synapse in LHb of parental separation model. (a) Representative images of ALDH1L1-RFP fluorescence and VGLUT2, and Homer1 staining in the LHb and quantification of ALDH1L1-RFP and Homer1 co-localisation, ALDH1L1-RFP and VGLUT2 co-localisation, VGLUT2 and Homer1 co-localisation in the Control and Separation groups (n = 5 mice, Scale bar, left, 5 μm, right, 1 μm). (b) Representative images of ALDH1L1-RFP fluorescence and Synapsin, and Gephyrin staining in LHb, and quantification of ALDH1L1-RFP and Synapsin co-localisation, ALDH1L1-RFP and Gephyrin co-localisation, and Synapsin and Gephyrin co-localisation in the Control and Separation groups (Control n = 3 mice, Separation n = 5 mice, Scale bar, left, 5 μm, right, 1 μm).

**Extended Fig 5.**
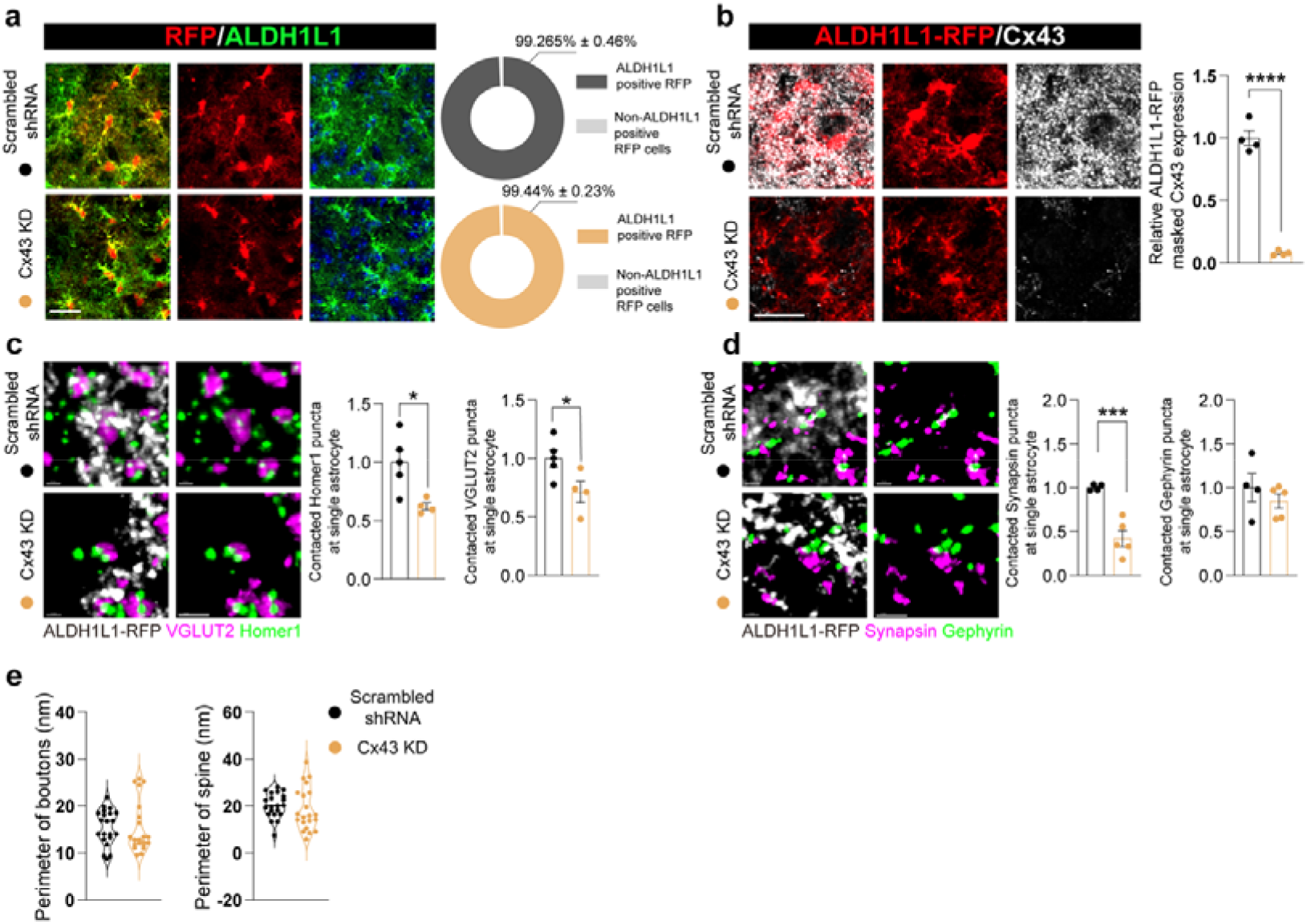
Efficacy and specificity of Cx43 knockdown by AAV and its effect on tripartite synapses. (a) Representative images of RFP fluorescence and ALDH1L1 staining in LHb by Cx43 knockdown (Cx43 KD) or scrambled control AAV injection (Scrambled shRNA), and percentage of ALDH1L1 positive RFP cells (n = 5 mice, Scale bar, 20 μm). (b) Representative images of ALDH1L1-RFP and Cx43 staining in the LHb and quantification of Cx43 expression in the Scrambled shRNA and Cx43 KD mice (n = 4 mice, Scale bar, 20 μm). (c) Representative images of ALDH1L1-RFP labelled astrocytes co-stained with VGLUT2 and Homer1, and the number of triple co-stained puncta, as well as VGLUT2 and Homer1 co-localisation in LHb (Scrambled shRNA, n = 5 mice, Cx43 KD, n = 4 mice, Scale bar, 1 μm). (d) Representative images of ALDH1L1-RFP labelled astrocytes co-stained with Synapsin, and Gephyrin, and the number of triple co-localisation puncta of these markers, as well as Synapsin and Gephyrin co-localisation (Scrambled shRNA, n = 4 mice, Cx43 KD, n = 5 mice, Scale bar, 1 μm). (e) Quantification of perimeter of boutons and spine observed from TEM.

**Extended Fig 6.**
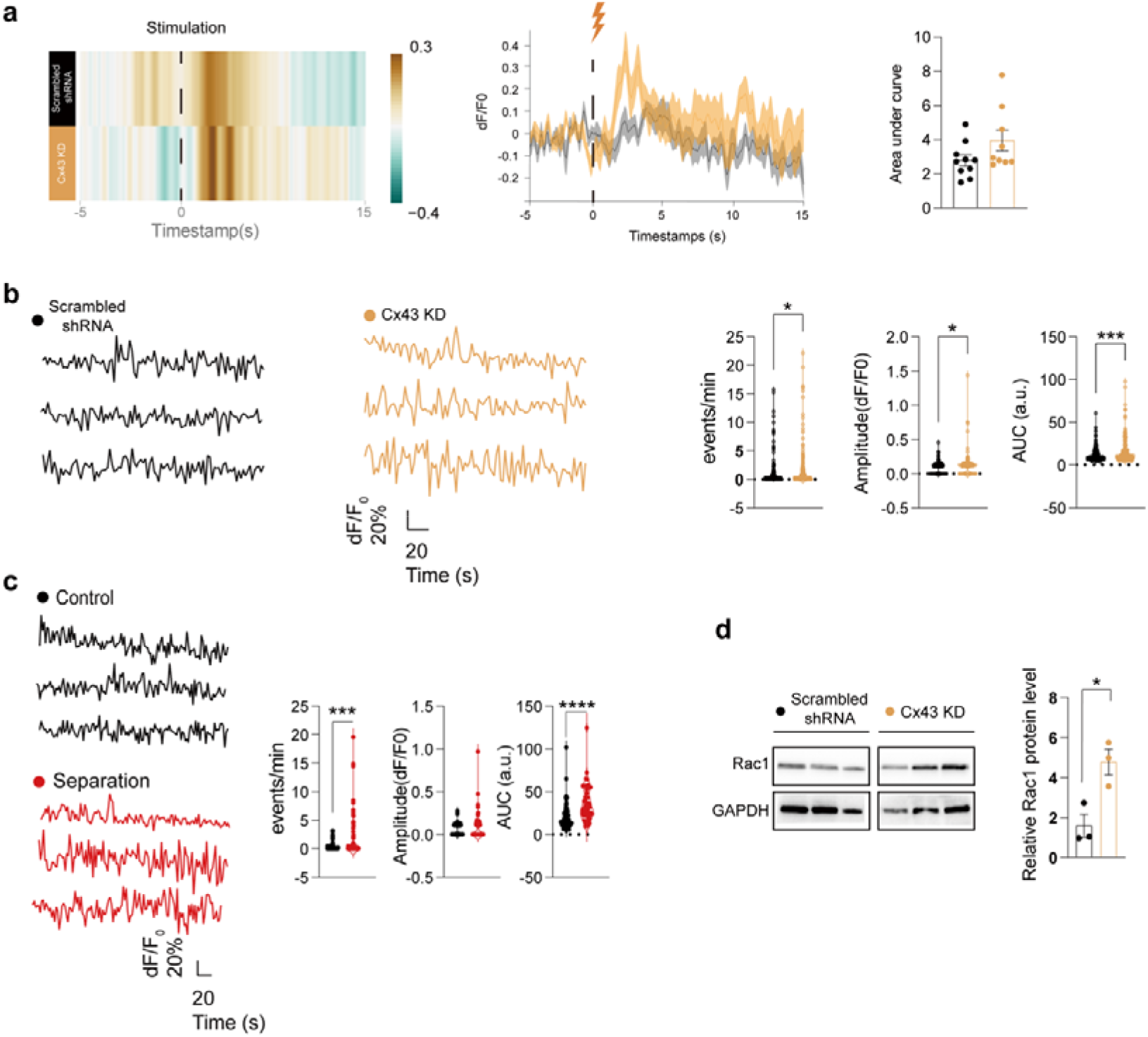
*Ex vivo* and *in vivo* recording of astrocyte Ca^2+^ signals. (a) Representative astrocytic [Ca^2+^]_i_ recordings from acute brain slices, and quantification of frequency, amplitude and area under the curve (AUC) of spontaneous Ca^2+^ signalling events in Control, Separation and Cx43 overexpression (Cx43 OE) group (Control n = 69 cells from 3 mice, Separation n = 61 cells from 3 mice, Cx43 OE n = 46 cells from 3 mice). (b) Representative astrocytic [Ca^2+^]_i_ recordings from acute brain slices, and quantification of frequency, amplitude and area under the curve (AUC) of Ca^2+^ signalling events in mice with Cx43 KD or Scrambled shRNA injection to LHb (Scrambled shRNA n = 192 cells from 6 mice, Cx43 KD n = 170 cells from 5 mice). (c) Heatmap and average curve of calcium signal assessed from *in vivo* fibre photometry recordings from Cx43 knockdown mice LHb astrocytes (n = 13 from 4 mice).

**Extended Fig 7.**
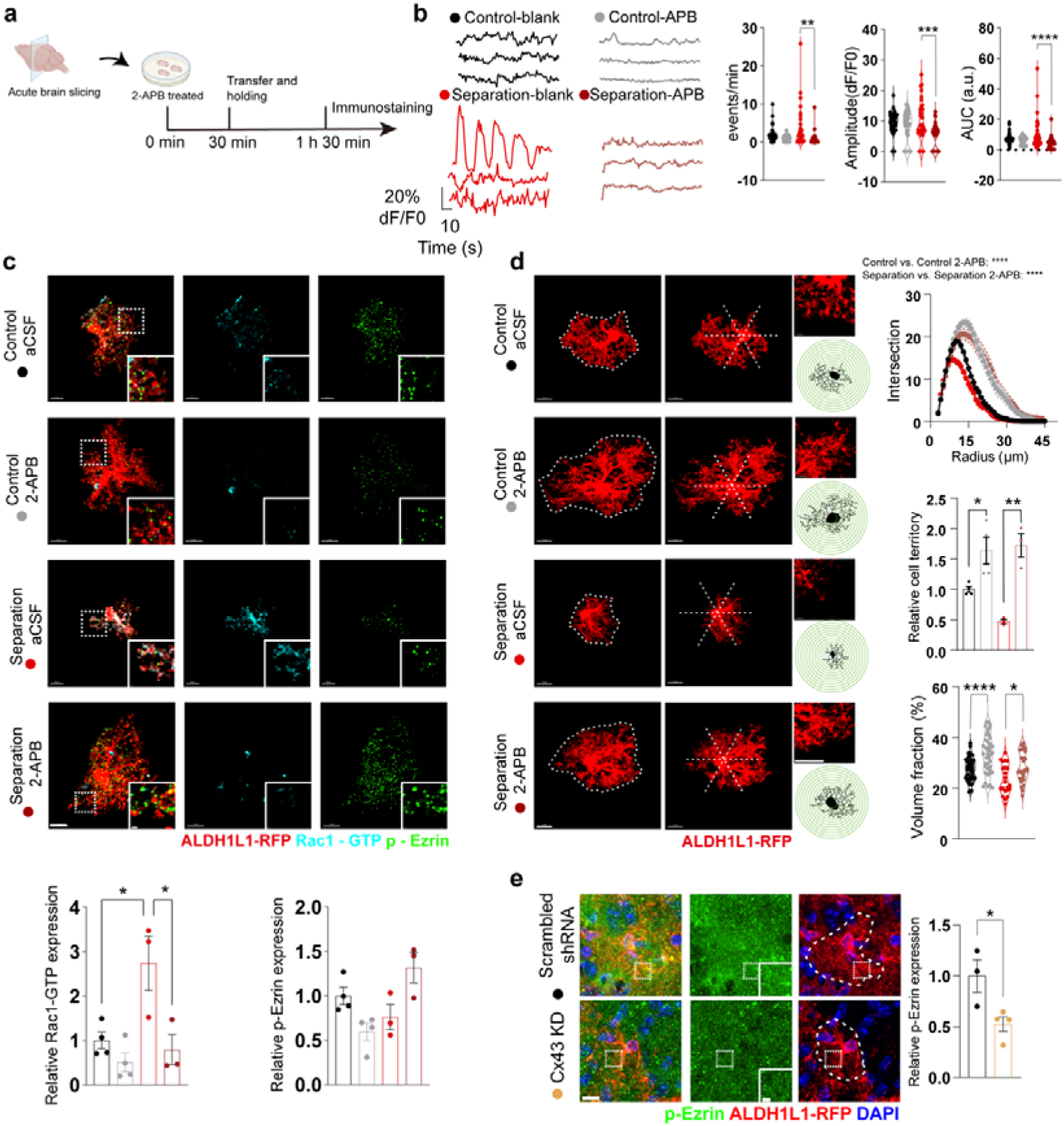
Intracellular Ca^2+^ signal mediated Rac1 expression of astrocytes is correlated with cell morphology in LHb. (a) Schematic paradigm of 2-APB application in acute brain slices. (b) Representative [Ca^2+^]_i_ recordings in control and 2-APB treated slices and Ca^2+^ dynamics quantification (Control aCSF, n = 36, Control 2-APB, n = 31, Separation aCSF, n = 36, Separation 2-APB, n = 42). (c) ALDH1L1-RFP labelled astrocytes co-stained with Rac1-GTP and p-Ezin, and relevant quantifications (Control, n = 4 mice, Separation, n = 3 mice; Scale bar: left, 10 μm, right, 1 μm). (d) Astrocytic territories (Control aCSF and Control 2-APB, n = 4 mice, Separation aCSF and Separation 2-APB, n = 3 mice), Sholl analysis (Control aCSF, n = 55 cells from 4 mice, Control 2-APB n = 48 cells from 4 mice, Separation, n = 33 cells from 4 mice, Separation 2-APB, n = 29 cells from 4 mice) and volume fraction (Control aCSF, n = 46 cells from 4 mice, Control 2-APB n = 42 cells from 4 mice, Separation, n = 13 cells from 3 mice, Separation 2-APB, n = 29 cells from 3 mice) of ALDH1L1-RFP labelled astrocytes in LHb of 2-APB treated slices (Scale bar: left, 10 μm, right, 5 μm). (e) Representative fluorescence images ALDH1L1-RFP-labelled astrocytes co-stained with p-Ezrin antibodies and respective quantification in the Cx43 Knockdown mice (Scrambled shRNA, n = 3 mice, Cx43 KD, n = 4 mice, Scale bar: left, 10 μm, right, 2 μm).

**Extended Fig 8.**
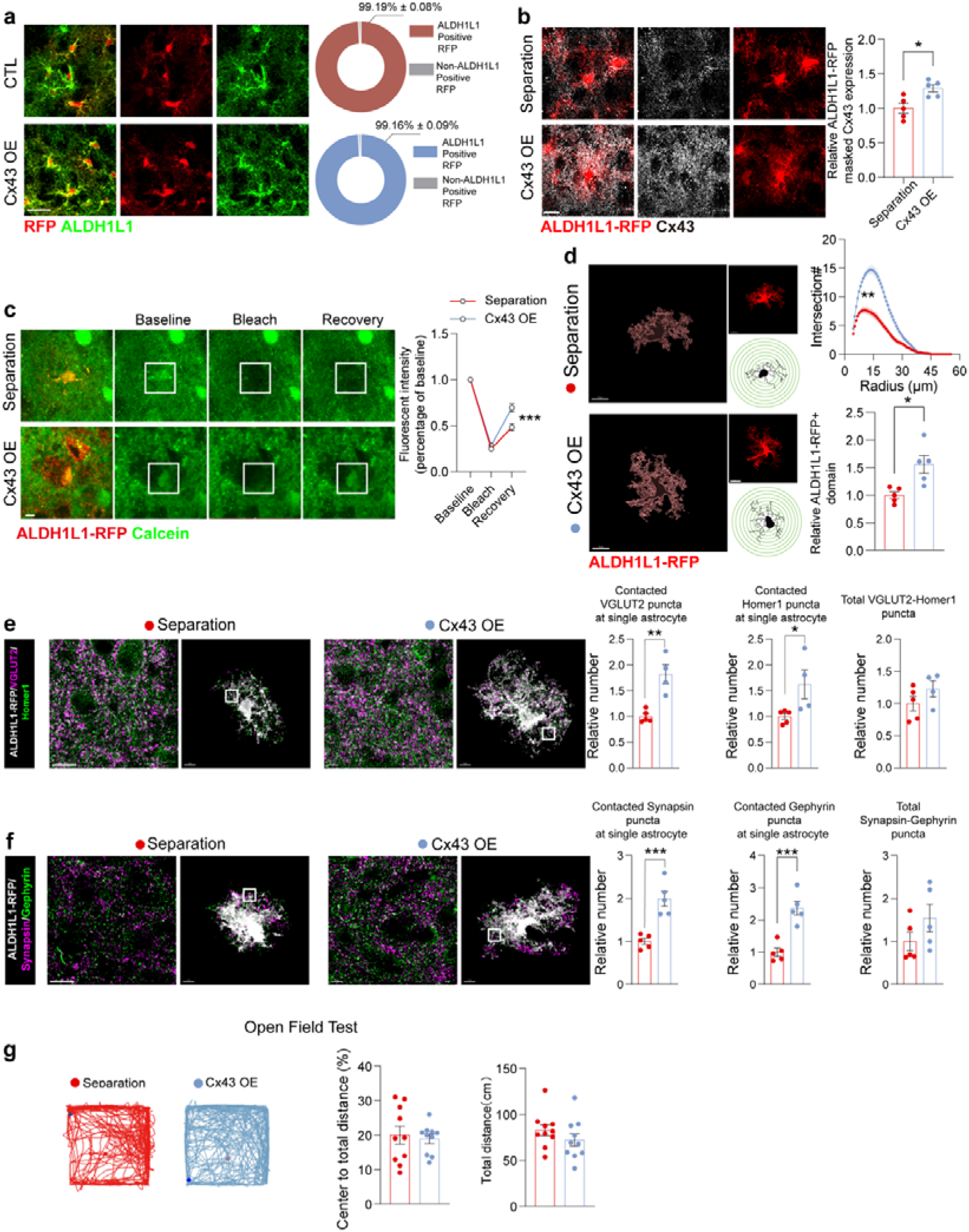
Efficacy and specificity of Cx43 overexpression by AAV, neuronal calcium activity and astrocyte-synaptic association in LHb after Cx43 overexpression. (a) Representative images of RFP and ALDH1L1 staining in the LHb after Cx43 overexpression (Cx43 OE) or scrambled control AAV injection (CTL), and percentage of ALDH1L1-positive RFP fluorescent cells (n = 5 mice, Scale bar, 10 μm). (b) Representative images of ALDH1L1-RFP labelling and Cx43 staining in the LHb and quantification of Cx43 positive area in the Separation and Cx43 OE mice (n = 5 mice, Scale bar, 10 μm). (c) Calcein based gap-FRAP assay and quantification of its fluorescence (n = 3 mice, Scale bar, 10 μm). (d) Representative fluorescence of astrocytic territories and skeletonised images of ALDH1L1-RFP labelled astrocytes, Sholl analysis and astrocytic domains (Sholl analysis, Separation, n = 82 cells from 3 mice, Cx43 OE, n = 77 cells from 3 mice, cell domains, n = 5 mice, Scale bar, 10 μm). (e) Representative images of ALDH1L1-RFP labelled astrocytes co-stained with VGLUT2 and Homer1, and quantifications of ALDH1L1-RFP and VGLUT2 co-localisation, ALDH1L1-RFP and Homer1 co-localisation, ALDH1L1-RFP and Homer1 co-localisation, VGLUT2 and Homer1 co-localisation (Separation n = 5, Cx43 OE n = 4). (f) Representative images of ALDH1L1-RFP labelled astrocytes co-stained with Synapsin and Gephyrin staining, and quantification of ALDH1L1-RFP and Synapsin co-localisation, ALDH1L1-RFP and Gephyrin co-localisation, Synapsin puncta and Gephyrin co-localisation (n = 5). (g) Open Field Test of the Separation group or Cx43 OE group mice (n = 10 mice).

## Notes

### Competing Interest Statement

The authors have declared no competing interest.

